# Continuous variable response, kinetic gating and connectivity that govern IS topology are perturbed in hyperinsulinemia

**DOI:** 10.1101/2020.08.10.243675

**Authors:** Namrata Shukla, Shantanu Kadam, Ranjith Padinhateeri, Ullas Kolthur-Seetharam

**Affiliations:** Department of Biological Sciences, Tata Institute of Fundamental Research, Mumbai, Maharashtra, 400005, India; Department of Biosciences and Bioengineering, Indian Institute of Technology Bombay, Mumbai, Maharashtra 400076, India

## Abstract

Understanding kinetic control of biological processes is as important as identifying components that constitute pathways. Insulin-signaling (IS) is central for almost all metazoans and its perturbations are associated with various diseases and aging. While temporal phosphorylation changes and kinetic constants have provided some insights, constant or variable parameters that establish and maintain signal topology are poorly understood. Our iterative experimental and mathematical simulation-based approaches reveal novel kinetic parameters that encode concentration and nutrient dependent information. Further, we find that pulsatile fasting insulin rewires IS akin to memory and in anticipation of a fed response. Importantly, selective kinetic gating of signals and maximum connectivity, between metabolic and growth-factor arms under normo-insulinemic states, maintains network topology. In addition to unraveling kinetic constraints that determine cascade architecture, our findings will help in identifying novel therapeutic strategies that conserve coupling between metabolic and growth-factor arms, which is lost in diseases and conditions of hyperinsulinemia.

## Introduction

Signaling cascades are essential for regulating cellular processes and decades of work has unraveled molecular and biochemical mechanisms that constitute them. However, kinetic parameters that define emergent properties of signaling networks and therefore predict regulatory nodes are poorly understood. While independent experimental and mathematical approaches have provided valuable insights (Behar et al., 2008; Faro et al., 2017; Kubota et al., 2012; Shinar et al., 2007; Somvanshi et al., 2019; Vinod and Venkatesh, 2009; Wilson et al., 2017), studies that capture dynamics and complexities of signaling architecture vis-à-vis physiological variations in input strengths are far fewer. Not only would these reveal fundamental kinetic considerations that determine signal topology but also inform about reactions/entities that could emerge as therapeutic targets.

Insulin signaling (IS), an evolutionarily conserved mechanism is essential for cellular/organismal metabolism and growth (Boucher et al., 2014; Haeusler et al., 2018; Saltiel and Kahn, 2001). Aberrant IS is associated both causally and consequentially with growth abnormalities, inflammation, accelerated aging and diseases including metabolic disorders and cancer (Arcidiacono et al., 2012; Guo, 2014; Hill and Milner, 1985; Shimobayashi et al., 2018; Shoelson et al., 2006; Vigneri et al., 2020). Genetic perturbations and omics-based studies have elucidated importance of key phosphorylation events in response to insulin stimulation (Humphrey et al., 2015; Krüger et al., 2008; Schmelzle et al., 2006; Yugi et al., 2014). Recent reports have provided crucial insights into physical protein interactomes, temporal changes in phospho-proteome and kinetic constants, viz T_1/2_ and EC_50_ (Kubota et al., 2018; Vinayagam et al., 2016). However, kinetic parameters that govern network properties of IS as a function of normo-insulinemic and hyper-insulinemic states that could collectively determine physiological and pathophysiological outcomes is still lacking.

Our current understanding largely stems from studies, which have used either supra-physiological or static concentrations of insulin. It is important to note that circulating insulin concentrations vary drastically from being low/pulsatile to high/biphasic in fasted and fed states respectively (Krishnan et al., 2018; Lu et al., 2012; Pørksen, 2002; Vander Haar et al., 2007). Moreover, hyper-insulinemia is associated with metabolic disorders such as diabetes and obesity (Menge et al., 2011; Satin et al., 2015; Schmelzle et al., 2006). These are key considerations since kinetic criteria that either encode fasted-to-fed transitions or drive pathological manifestations of IS are unknown. Furthermore, IS can be broadly divided into metabolic and growth factor arms (Mendoza et al., 2011; Petersen and Shulman, 2018). In this regard, while biased signaling is implicated in diseases, if/how the flow of information is stratified and maintained remains to be unraveled.

Mathematical approaches to model cellular signaling have gained traction in the recent past to understand the dynamics and also to provide predictive parameters that define topology of signaling network (Cedersund et al., 2008; Dalle Pezze et al., 2016; Di Camillo et al., 2016; Sedaghat et al., 2002; Sonntag et al., 2012). Earlier such attempts to determine kinetics of insulin signaling have largely employed “averaged” measures to define the behavior of the system (Kubota et al., 2018). Importantly, given the fluctuations in insulin levels and inherent noise in signaling, there are no reports that have computed kinetic parameters, which capture emergent properties of IS. Specifically, while there have been simulation based approaches to define dose-to-duration effects and kinetic insulation on synthetic signaling networks (Behar et al., 2007), such principles have not been applied to complex cascades such as insulin signaling.

In this regard, our current study addresses how connectedness among signaling components as well as overall network topology is maintained under physiological concentrations of insulin. We further highlight the concentration dependency of barriers in the signaling cascade which maintain hierarchy. Additionally, our study puts emphasis on the importance of dynamic range and pulsatility in signaling which generates memory as well as couples the metabolic and growth factor arms.

## Results

### Distinct kinetics of signaling in response to physiological and non-physiological concentrations of insulin

Although previous reports have attempted to elucidate dynamics of insulin signaling, kinetic parameters that define signaling architecture in response to physiologically relevant insulin concentrations remain to be unraveled. This is particularly important since circulating concentrations of insulin vary between 0.1 nM and 1.0 nM during normal fed-fast cycles. Moreover, insulin signaling achieves both nutrient uptake and its utilization via anabolic processes, whose perturbations are associated with diseases and accelerated aging. Thus, we wanted to assess the kinetics of signaling through nodal kinases in the cascade, which govern both the metabolic and growth factor arms (Figure 1A). Given the importance of liver in modulation of insulin action and integration of whole-body physiology, we employed primary hepatocytes. Towards this, primary hepatocytes isolated from mice livers were treated with different concentrations of insulin as described in Figure 1B. Our paradigm ensured that the kinetic evaluation did not have any bearing from either residual signals or nutrient inputs alone, as illustrated in Figure 1B and Figure 1- figure supplement 1, A-D.

**Figure 1:**
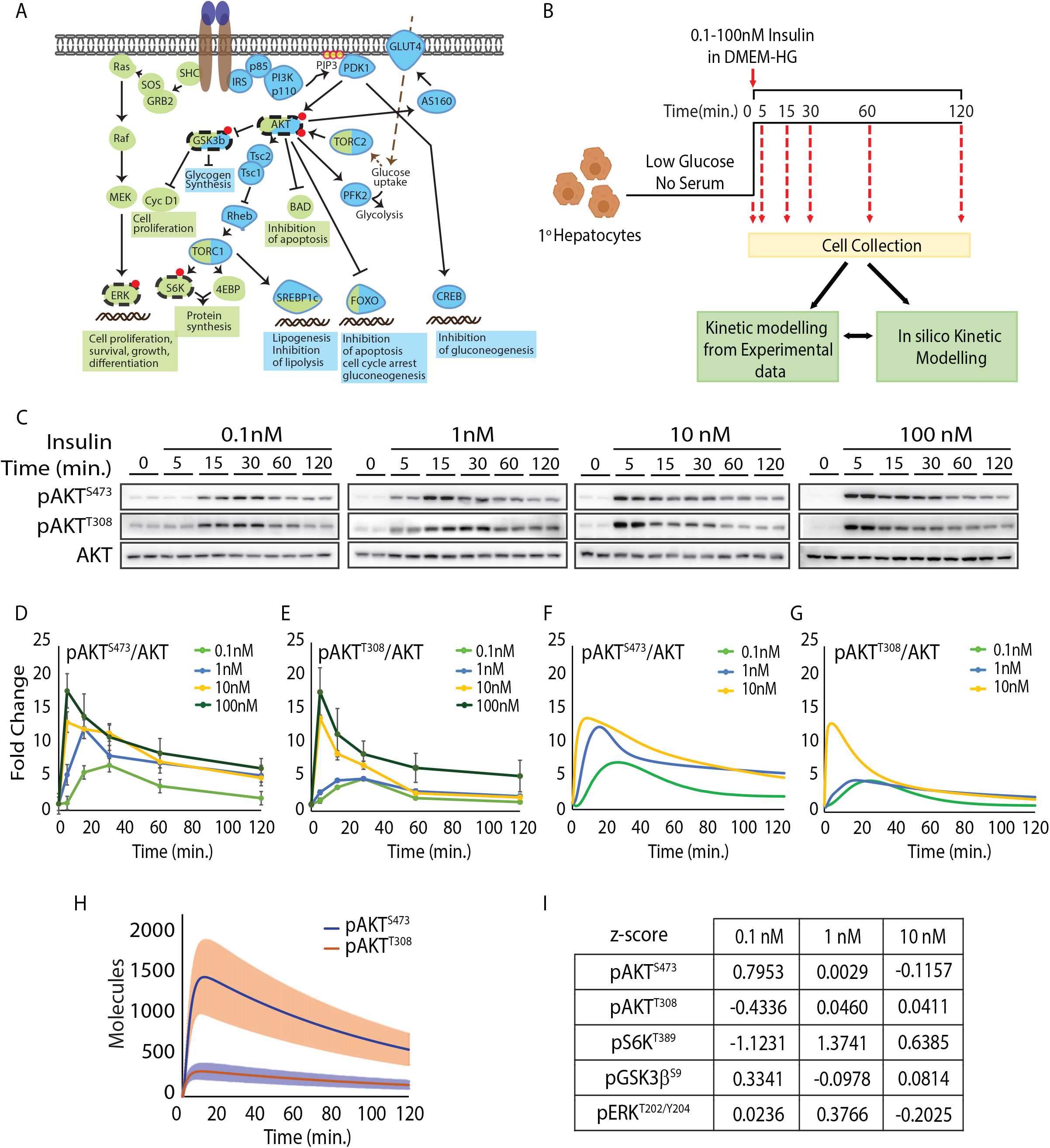
Iterative experimental-mathematical approach reveals distinct insulin signaling kinetics. (A) Schematic of the insulin/IGF signaling pathway. Components involved in metabolic and mitogenic arms are shown in blue and green, respectively. Phosphorylations measured in this study are highlighted in red. (B) Experimental paradigm and workflow for assaying signaling in response to one step stimulation. (C) Representative blots for levels of pAKT^T308^ and pAKT^S473^ following insulin stimulation, as indicated. Total AKT and actin were used for normalization. See more in Figure 1- figure supplement 1E-F and 2A-B. (D) and (E) Quantitation for temporal changes in pAKT^S473^ (D) and pAKT^T308^ (E) from experimental data shown in C. Fold changes for each concentration are with respective to their own 0m time point. Data presented is mean ± s.e.m. (N=4, n=4). (F) and (G) Quantitation for temporal changes in pAKT^S473^ (F) and pAKT^T308^ (G) from mathematical simulations using differential equations. (H) Kinetic behavior of phosphorylated pAKT^T308^ and pAKT^S473^ molecules at 1 nM insulin from stochastic simulations. The band represents standard deviation. (I) z-score giving degree of concordance between simulated and experimental data.

As reported by others (Borisov et al., 2009; Kubota et al., 2012; Kubota et al., 2018; Noguchi et al., 2013) insulin treatment led to a rapid activation of downstream signaling and was consistent with a fed response (Figure 1C-E, Figure 1- figure supplement 1E-F and Figure 1- figure supplement 2A-B). Expectedly, overall signal intensities (i.e. area under the curve: AUC), for all the phosphorylation events scored in our assay, were positively correlated with insulin concentration (Figure 1- figure supplement 2C). We define true discovery rate as a statistical measure to compare changes in phosphorylations across time and insulin concentrations. This parameter further validates the statistical robustness of our measurements (Figure 1- figure supplement 2D). It was striking to see that the kinetic behavior of nodal kinases in the cascade, AKT and ERK, was markedly different (Figure 1D-E, Figure 1- figure supplement 1E-F, 2A-B and 3C), which has not been highlighted in any of the previous studies. Importantly, we observe non-linear and non-monotonic association of signal intensities w.r.t insulin concentrations, across the cascade, both in terms of extent of phosphorylation and temporal behavior.

For example, activation-inactivation kinetics was starkly different for AKT (T^308^ and S^473^) and ERK. In addition to this, while the final intensity of AKT phosphorylation approached baseline by 120 minutes, ERK phosphorylation showed a distinct second wave of activation (Figure 1D-E, Figure 1- figure supplement 1E-F, 2A-B and 3C). Similarly, we found that initial induction of phosphorylation of GSK3β and S6K was phase delayed in response to 0.1 nM and 1.0 nM insulin treatments and continued to remain elevated long after phosphorylation on AKT started to extinguish (Figure 1- figure supplement 1E-F, 2A-B and 3A-B).

Further, on comparing both fed and fasted insulin concentrations, it was apparent that maximal phosphorylation and its sustenance varied drastically for nodal kinase AKT, as can be seen in Figure 1D-E. These clearly indicated that insulin-dependent programming of signaling kinetics was distinct and prompted us to investigate the kinetic parameters that defined this behavior. We adopted an iterative experimental-cum-mathematical approach to gain further insights.

### Mathematical modelling of signaling kinetics and predictive assessment of key phosphorylation-dephosphorylation dynamics

Based on our experimental results, we modeled the signaling cascade using mathematical methods with an aim to extract kinetic parameters that define the network. We considered the insulin signaling network as a set of coupled biochemical reactions and used ordinary differential equations (ODE) to describe the system (Aoki et al., 2013; Arkun, 2016; Borisov et al., 2009; Dalle Pezze et al., 2016; Dalle Pezze et al., 2012; Di Camillo et al., 2016; Ho et al., 2015; Huang et al., 2014; Kubota et al., 2012; Kubota et al., 2018; Noguchi et al., 2013; Sedaghat et al., 2002; Zhao et al., 2017). Importantly, we set out to not only test the robustness of our mathematical simulation using experimental results, but also predict the behavior of components that were not measured experimentally.

As shown in Figure 1F-1G and Figure 1 – figure supplement 3D-F, the simulation results for pAKT^T308^, pAKT^S73^, pGSK3β^S9^, pS6K^T389^ and pERK^T202/Y204^ were consistent and nearly overlapping with the experimental data, across insulin concentrations. Next, using experimentally optimized parameters as input, we simulated the phosphorylation dynamics of insulin receptor (IR), mTORC1 and mTORC2 (Figure 1 – figure supplement 3G).

While downstream components scaled with insulin concentration, the most upstream event of insulin receptor phosphorylation was rapid and transient. Intriguingly, kinetics of mTORC1 and mTORC2 were starkly different (Figure 1 – figure supplement 3G). Moreover, even though mTORC2 is not directly downstream to IR, we found their responsivity to be similar qualitatively. To our best knowledge this is one of the first attempts that delineates temporal variations in activation of mTOR complexes. Given that mTORC2 is the primary kinase for S^473^ phosphorylation, the discordant dynamics of mTORC2 phosphorylation and pAKT^S473^ predicts additional regulatory steps in controlling activation of AKT (Figure 1D and Figure 1 – figure supplement 3G).

We also used stochastic simulations to provide an alternative approach to validate our mathematical predictions, which qualitatively resembled the deterministic approach for quantities as described in Figure 1H. In addition to predicting the response at a population level, this allowed us to determine the fluctuations in the system and compare it with signaling topology (see below).

In order to score the robustness of our simulation data, we computed the z-score for all components assessed in the signaling cascade across concentrations (Figure 1I). It is important to note that our iterative toggling between computational and experimental determination of phosphorylation gave highly consistent results. Further, low values of false discovery rate calculations suggested statistical similarity between simulation and experimental data (Figure 1 – figure supplement 3H).

### AKT-dependent responsivity to insulin is determined by phosphorylation at S^473^

Several reports have highlighted the necessity of dual phosphorylation of AKT at T^308^ and S^473^ for its activity (Bertuzzi et al., 2016; Manning and Toker, 2017; Sarbassov et al., 2005). Despite this it is still unclear as to which of these provides the gain in terms of signal strength and responds to dynamic changes in insulin concentrations. Thus, we computed percentage gain in signal against physiological insulin concentrations of 0.1 and 1 nM for pAKT^T308^ and pAKT^S473^ (Figure 2A). pAKT^T308^, that is directly downstream to insulin receptor, showed comparable activation with no change in peak intensity across fasted and fed insulin concentrations. On the contrary, pAKT^S473^, which is indirectly dependent on insulin via mTORC2, displayed dose responsiveness to insulin concentrations and variable kinetics (Figure 2A-B). This finding posits that while pAKT^T308^ may serve to prime the signaling, pAKT^S473^ determines the extent of overall activation in response to fed insulin doses.

**Figure 2:**
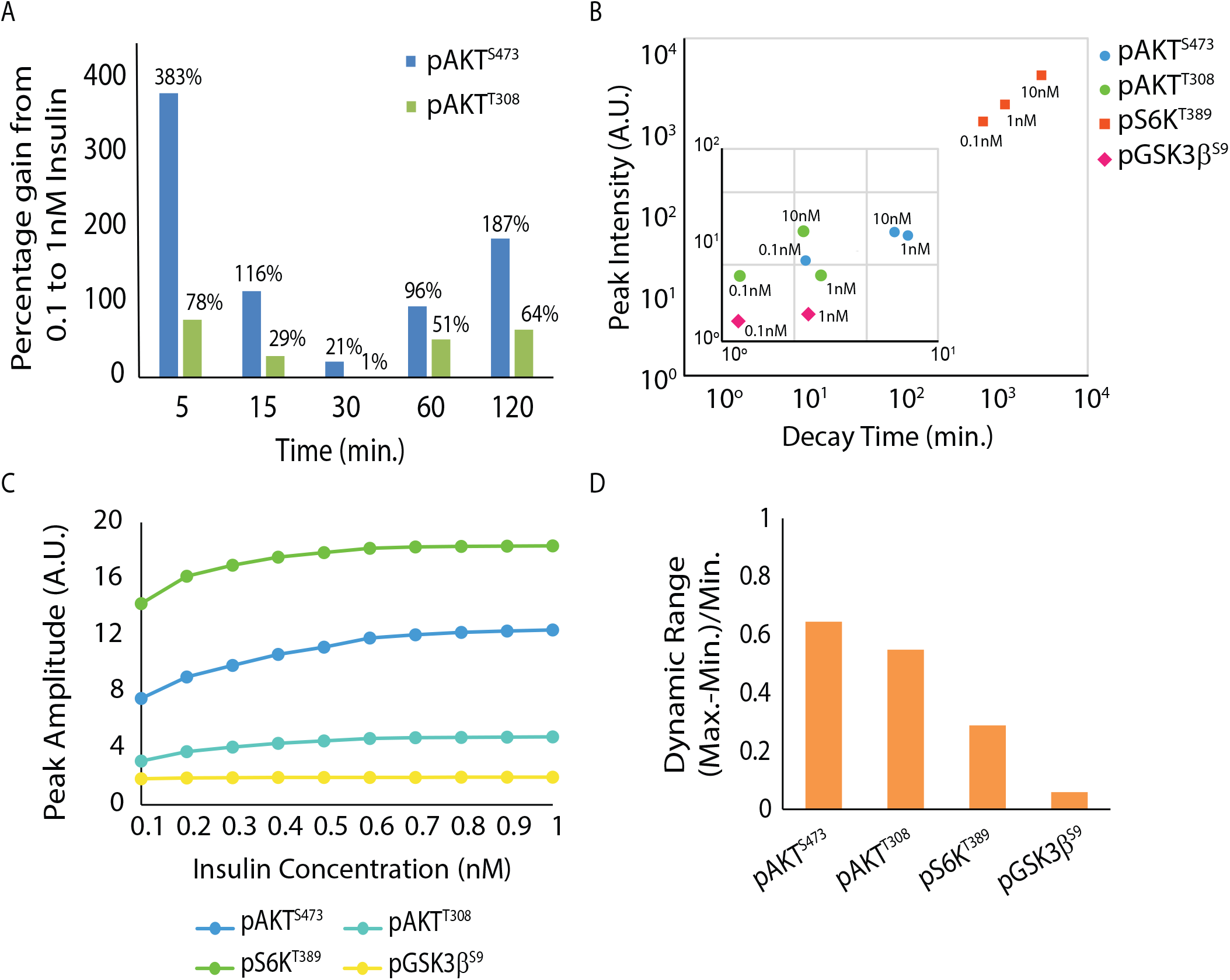
Continuous variable parameters and high/low pass filters determine concentration dependent insulin response. (A) Extent of change in phosphorylation at pAKT^T308^ and pAKT^S473^ across time points between 0.1 and 1 nM insulin. (B) Phase diagram depicting relationship between peak intensity and decay time from simulated data. (C) Estimated peak amplitude for signaling components as a function of varying insulin concentrations depicts dynamic range. (D) Dynamic range for signaling components computed from (C), as labeled.

### Non-concordant peak and final amplitudes define dynamic range of nodal signaling events

Threshold of activation and dynamic range are key determinants of responsivity in signaling, especially when the inputs are dynamic, as in the case of IS. We wanted to determine (a) the relationship between peak intensity and decay kinetics, and (b) dynamic range and threshold activation, which collectively dictate the physiological output. Phase diagram depicting peak amplitudes and decay time of phosphorylation events highlighted non-concordance between these for AKT but not for pGSK3β and pS6K (Figure 2B).

Most experimental approaches in the past have assayed for signaling in response to very high inputs, which is rarely physiological. Such deterministic evaluation of signaling also masks threshold kinetics, which is critical to encode biological response. Therefore, on simulating the phosphorylation events across concentrations from 0.1 nM to 1.0 nM, we found disparate dynamic ranges for activation (Figure 2C). While AKT phosphorylations (both at S^473^ and T^308^) displayed large and nearly overlapping dynamic range, pS6K^T389^ and pGSK3β^S9^ reach saturation at lower concentrations of insulin (Figure 2C-D). Taken together with dose-dependency of pAKT^S473^ (Figure 2A), these results clearly suggest that while dual phosphorylation of AKT is important for its activity, pAKT^S473^ is a crucial regulatory node during fast to fed transitions.

Since our simulations predicted non-saturation dynamics for pAKT^S473^, we wanted to experimentally verify if this was indeed the case. We specifically chose 0.3 nM and 0.6 nM, as the response is linear at 0.3 nM and begins to plateau at 0.6 nM insulin. As shown in Figure 2 – figure supplement 1A-B, our experimental results were consistent with the mathematical predictions and clearly indicated that pAKT^S473^ indeed displayed a large dynamic range to insulin inputs.

Interestingly, the variability in dynamic range was independent of the final amplitudes (Figure 2 – figure supplement 1C) as it returned to the same level at 120min, for all the phosphorylations assessed. Such non-concordance between peak and final amplitudes across signaling components raised the exciting possibility of existence of (a) kinetic insulation of signals and (b) memory of fasted insulin inputs, which together would define the fed insulin response.

### Diverse insulin inputs generate differential kinetic gates and signal noise

In a multi-component and multi-step signaling cascade, such as insulin signaling, it is important to determine parameters that (a) define the topology or the information flow through the network and (b) those that maintain robustness of the network/topology. Kinetic insulation has been proposed as one of the key determinants of non-uniform flow of information. Although inferred by mathematical approaches (Behar et al., 2007; Behar et al., 2008) on a synthetic cellular signaling cascade, it has not been applied to a dynamic physiological system such as insulin signaling.

In this context, we used our experimental data and mathematical simulations (methods) to deduce kinetic gates that define topology of insulin signaling. To reveal kinetic gating/insulation we computed rate constants for phosphorylation events, which included known feed-forward and feed-back regulatory inputs (Figure 3- figure supplementary 1A-B). A simple-minded assumption was that very high or low ratios of K_ON_/K_OFF_ would constitute kinetic “gates” that determined differential flow of signals. A phase diagram of K_ON_/K_OFF_ ratios for key phosphorylation events is depicted in Figure 3A, wherein we applied 10^+/-1^ as the threshold or barrier.

**Figure 3:**
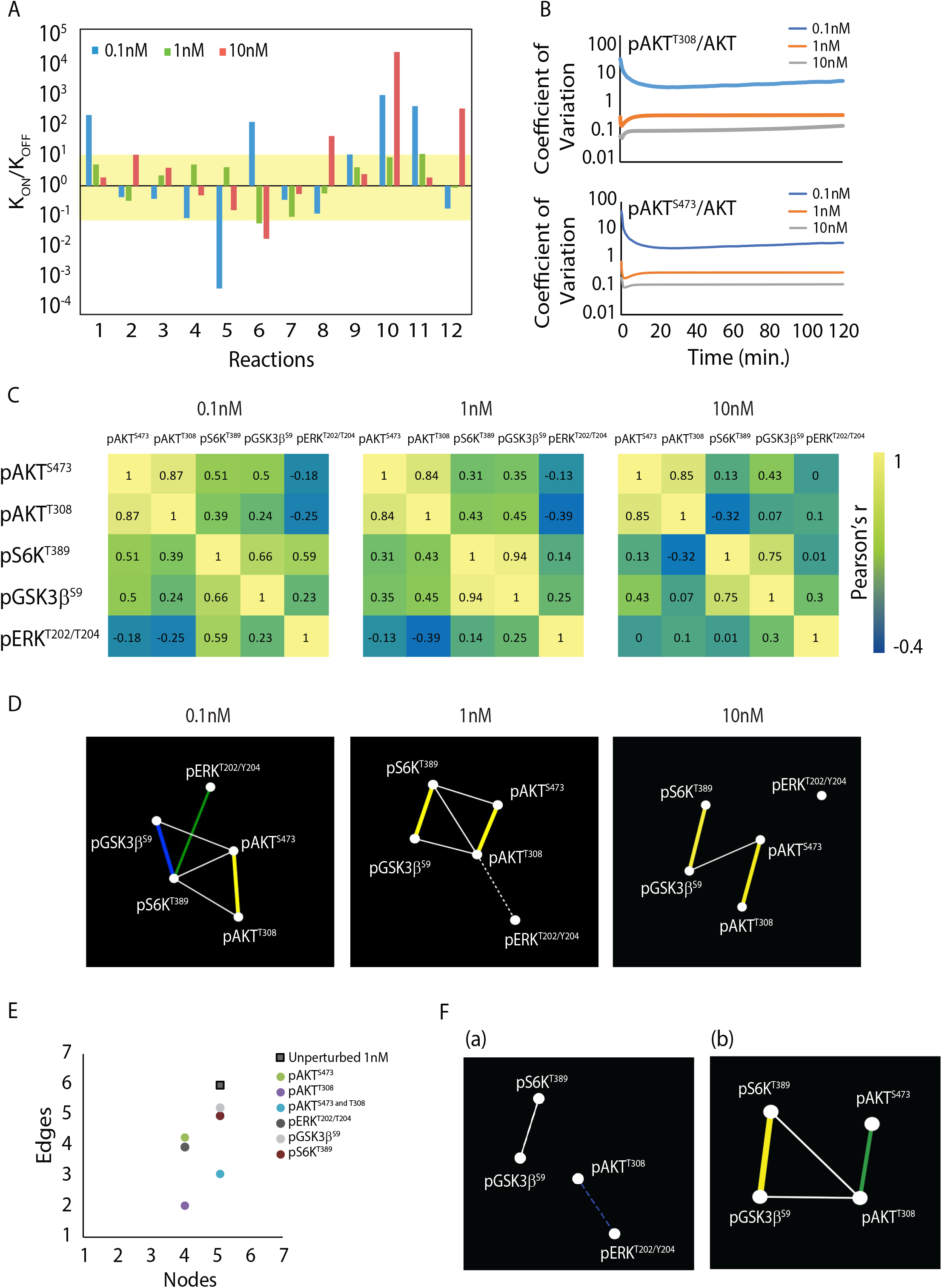
Kinetic gates and maximum connectedness is associated with robust topology under normo-insulinemic states. (A) K_ON_/K_OFF_ ratios for phosphorylations across the signaling cascade. Insulin concentrations of 0.1-10 nM are depicted separately. Numbers on the x-axis represent phosphorylation events as detailed in Figure 3 – figure supplement 1A. Yellow band refers to the kinetic barrier applied between 0.1-10, representing a 10-fold change. (B) Noise in signal for phosphorylation at pAKT^T308^ and pAKT^S473^ in response to different insulin concentrations, as indicated. (C) Correlation matrix depicts degree of relatedness between phosphorylation events and their evolution with increasing insulin concentration. (D) Network analysis depicting degree of connectedness across insulin concentrations, as indicated. Dashed line represents negative correlation. Significance in correlation: White (p<0.05) <Blue (p<0.005) <Green (p<0.0005) <Yellow (p<0.00001) as observed by Student’s t-test. (E) Number of edges and nodes in a 1 nM network substituted with 10 nM values, as indicated. (F) Network maps of 1 nM insulin perturbed with 10 nM pAKT^T308^ (a) and pAKT^S473^ (b)

Interestingly, at insulin concentration of 1 nM, which mimics a physiologically fed state, most reactions were not gated and were unlike the response to very low and very high insulin concentrations. For example, AKT activation (reaction 5) was more sensitive at lower insulin concentration i.e. there was a negative barrier while at 1 and 10 nM there was no gating. On the contrary, activation of GSK3β (reaction 10) was highly gated at both very low and very high insulin concentrations. We also observed a peculiar pattern between nodal priming events {viz. reactions 1 (Ins+IR ⇄ p1IRC), 6 (ppAKT+mTORC1 ⇄ ppAKT+pmTORC1), 10 (ppAKT+GSK3β ⇄ ppAKT+pGSK3β) and 11 (p1IRC+Raf ⇄ p1IRC+Raf*)} and their effector or downstream phosphorylations {reactions 4 (p1IRC+pAKT^S473^ ⇄ p1IRC+ppAKT), 5 (pmTORC2+pAKT^T308^ ⇄ pmTORC2+ppAKT) and 12 (Raf*+ERK ⇄ Raf*+ppERK)} vis-à-vis kinetic gates specifically at 0.1 nM. Together, it was striking to see that strong negative and positive barriers were differentially associated with metabolic and growth factor arms of the cascade in response to fasted, fed and supra-physiological insulin inputs.

Since phosphorylation of AKT is one of the central events that is used as surrogate for IS, and given the differential dynamics of pAKT^T308^ and pAKT^S473^, we wanted to assess their individual contributions to functional flexibility. We went ahead to compute noise in their signaling (fluctuations on mathematically determined concentrations; see methods). This is relevant as often noise in biology becomes important for mounting a robust response in addition to generating functional heterogeneity and flexibility especially in a dynamic system like IS (Bowsher et al., 2013; Silva-Rocha and de Lorenzo, 2010; Thattai and Van Oudenaarden, 2001). As shown in Figure 3B, we observed that lower concentrations of insulin generate more noise than higher concentration across time points assessed. While being in general agreement with similar measurements of other biological parameters, this also indicated that the differential phosphorylation dynamics of AKT^T308^ and AKT^S473^ were independent of noise. In summary, the results described in this section clearly indicated that differential insulin inputs mounted diverse kinetic responses, which together could possibly exert a control over topology of the cascade.

### Robust IS topology is achieved at physiological insulin inputs

Topology and robustness of a network is governed by the degree of connectedness among the network components and is defined by how correlated their responses are. Therefore, we set out to ask if supra-/physiological inputs of insulin had any bearing on signaling topology.

Computing Pearson coefficient across time for different insulin concentrations gave us a correlation matrix comparing each phosphorylation event with the other (Figure 3C). We observed that maximal correlations are lost under supraphysiological concentration of 10 nM compared to physiological concentrations of insulin.

Next, we checked if high degree of correlation in response to fed and fasted insulin inputs had any impact on the topology of the network. For a maximally connected network of “n” nodes the maximum number of edges would be n(n-1)/2; while the minimum number of edges would be (n-1). Applying this to a five-component system (as in our case) should give a maximum of 10 connections, although 34 undirected non-isomorphic graphs can be realized. We found that when 5 nodes (vertices) corresponding to pAKT^S473^, pAKT^T308^, pS6K^T389^, pGSK3β^S9^, pERK^T202/Y204^ were used, maximum connectivity was obtained at physiological concentrations of insulin (at 0.1 nM and 1.0 nM) (Figure 3D and Figure 3 - figure supplement 1C). Distinctively the network broke at 10 nM insulin and the node corresponding to pERK was disconnected, which indicated decoupling of the metabolic and growth factor arms with possible pathophysiological implications.

To understand which of the nodes control topology of the network, we substituted individual nodes of 0.1 and 1 nM insulin network with that of 10 nM while keeping the rest unperturbed. Perturbation of every component changed network properties with a reduction in both the number of nodes as well as edges (Figure 3E and Figure 3- figure supplement 2 and 3A-D). Interestingly, while perturbation of pAKT^S473^ caused disappearance of some edges, perturbation of pAKT^T308^ completely broke the network, bringing the connections down from 6 to 2 (Figure 3E and F).

### Pulsatile fasting insulin rewires response to fed insulin inputs akin to memory

Uniquely, insulin is released in a pulsatile manner during a fasted state (O’Meara et al., 1993; O’Rahilly et al., 1988), which is followed by a biphasic secretion in response to fed nutrient inputs. As mentioned earlier, while most studies on signaling dynamics have used high concentrations of insulin, there are no reports that have investigated kinetics and topology vis-à-vis pulsatile fasted insulin inputs. Moreover, if/how a fasted input shapes signaling architecture in a fed state has not been addressed, thus far.

To this end, we pulsed hepatocytes with 0.1 nM insulin and then subsequently treated with 1 nM insulin as a proxy to physiological dynamics of fasted and fed insulin inputs, as indicated (Figure 4A). Surprisingly, we found that there was neither sustenance nor an enhanced response to consequent insulin pulses, for pAKT^T308 and S473^ (Figure 4B-C and Figure 4 – figure supplement A), which was unanticipated. This striking loss of pAKT signal by the end of 4^th^ pulse (at 0’) was distinct from a continuous step treatment as described earlier (Figure 1D-E) and indicated a memory of signaling. This behavior was not seen for pERK (Figure 4D and Figure 4 – figure supplement B). Interestingly, pAKT levels reached a new baseline following pulsatile insulin stimulation (Figure 4B-C). This new reset point of pAKT also changed the kinetics following 1 nM insulin treatment, which was distinct from pERK, pGSK3β and pS6K phosphorylation (Figure 4D and Figure 4- figure supplement 1B-F). These results clearly indicated that fasted insulin pulses created a memory to possibly enhance the response to fed insulin inputs. In support of this hypothesis, network analyses of this pulsatile adapted fed IS showed more connectedness (as compared to 1 nM alone) (Figure 4E and Figure 4 – figure supplement 1G). Additionally, we also looked at the transcription of genes downstream of a pulsatile adapted system. In line with the signaling data, the transcription of target genes was also more robust post adaptation (Figure 4F).

**Figure 4:**
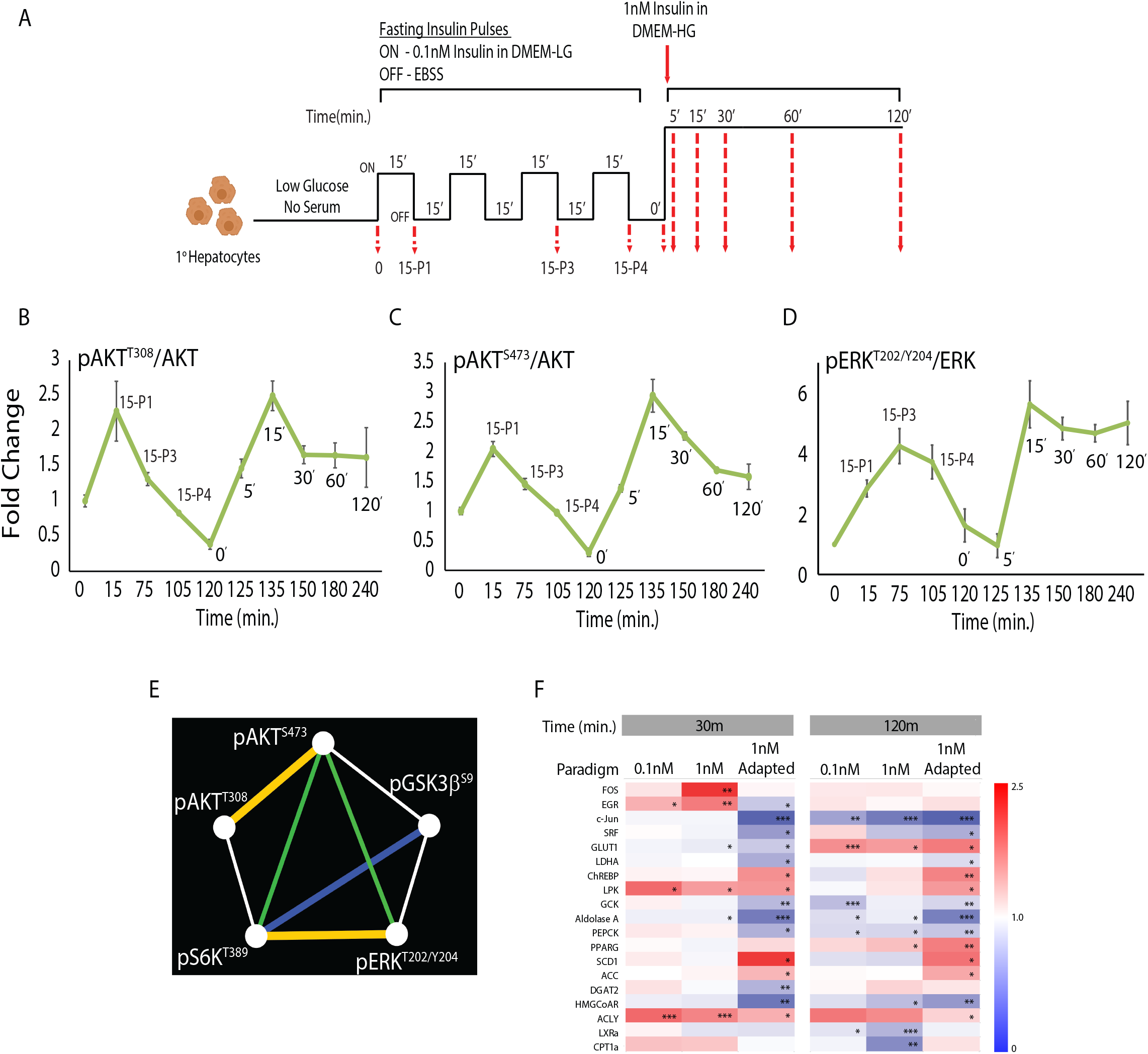
Pulsatile fasting insulin rewires response to fed insulin inputs akin to memory. (A) Experimental paradigm for mimicking fasted and fed insulin stimulation. P1-P4 indicate 0.1 nM insulin pulses. (B-D) Quantitation for temporal changes in phosphorylations at pAKT^T308^ (B), pAKT^S473^ (C) and pERK^T202/Y204^ (D) following insulin treatment as in A. Fold changes for each concentration are with respective to their own 0m time point. Data presented is mean ± s.e.m. (N=4). (E) Network analysis showing connectivity among signaling components after treatment with 1 nM insulin following fasted insulin inputs, as in A. Dashed line represents negative correlation. Significance in correlation: White (p<0.05) <Blue (p<0.005) <Green (p<0.0005) <Yellow (p<0.00001) as observed by Student’s t-test. (F) Heat maps for changes in gene expression downstream to insulin signaling in response to constant 0.1 nM and 1 nM, and 1 nM insulin following fasted insulin inputs (1 nM adapted, as in A) (N=2, n=3)

### Repeated stimulation by fed insulin abrogates the synergy between the metabolic and mitogenic arms of signaling

Although continuous exposure to higher levels of circulating insulin is known to cause resistance and thus metabolic diseases, the kinetic basis for such a signalling has not been investigated. In this context, we repeat stimulated hepatocytes with 1 nM insulin, as indicated in Figure 5A. This led to an anomalous response vis-à-vis both metabolic (pAKT) and mitogenic (pERK) arms of signalling. While pAKT levels decreased drastically, amplitude of pERK peaks increased following repeated stimulation (RS1 and 2) of fed insulin inputs (Figure 5B-D and Figure 5- figure supplement 1A). Network analysis following this paradigm showed complete loss of connections among signalling components (Figure 5E and Figure 5- figure supplement 1D). This was also apparent with the dynamics of pGSK3b and pS6K, which remain upregulated despite a downregulation in AKT signalling (Figure 5- figure supplement 1A-C). Interestingly, repeated stimulation of fed insulin also led to a loss in transcriptional robustness (Figure 5F).

**Figure 5:**
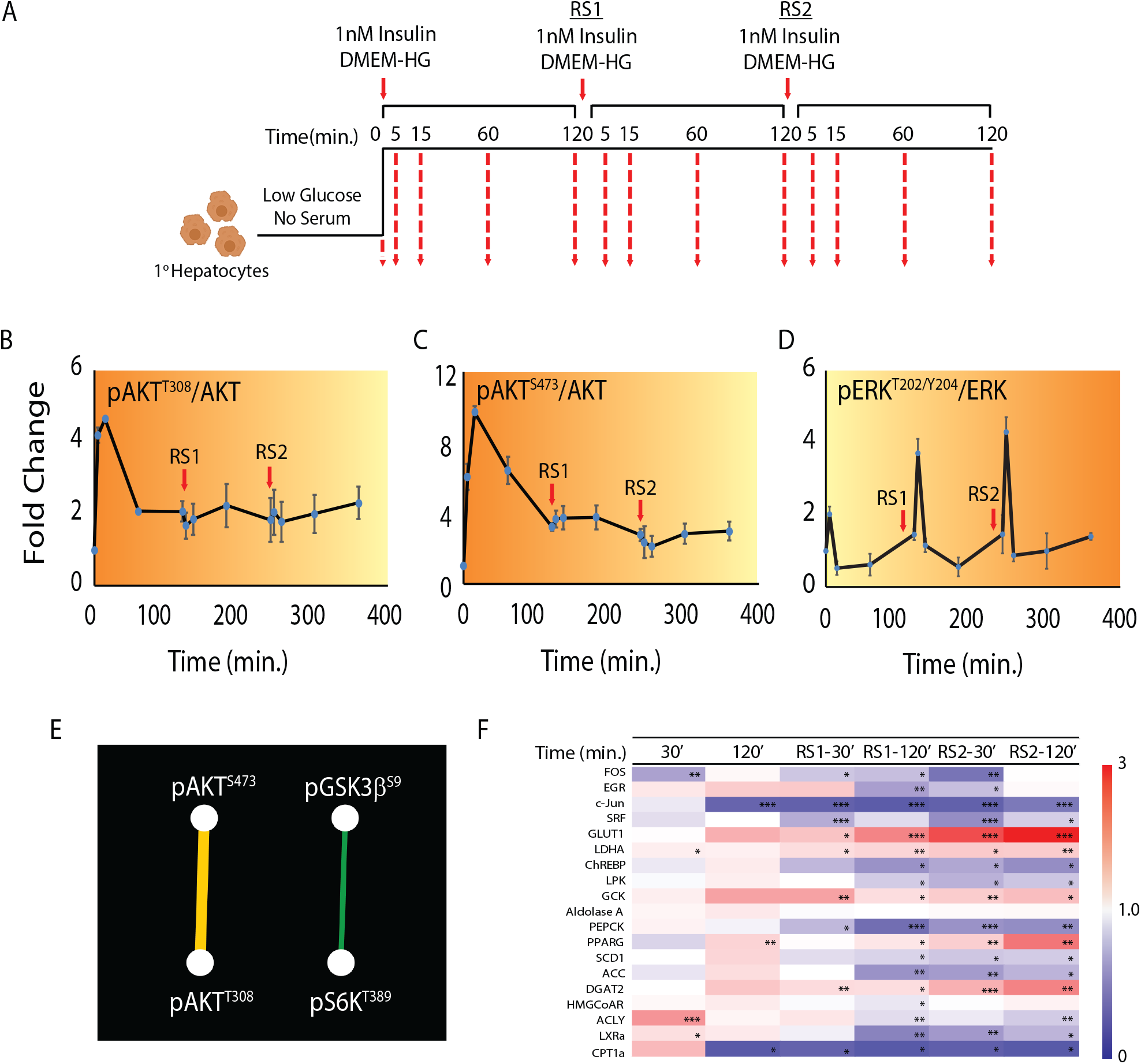
Repeated stimulation by fed insulin abrogates the synergy between the metabolic and mitogenic arms of signaling. (A) Experimental paradigm for repeated stimulation with 1nM insulin. (B-D) Quantitation for temporal changes in phosphorylations at pAKT^T308^ (B), pAKT^S473^ (C) and pERK^T202/Y204^ (D) following repeated insulin stimulation. Fold changes for each concentration are with respective to their own 0m time point. Data presented is mean ± s.e.m. (N=4). (E) Network analysis showing connectivity among signaling components after repeated insulin stimulation, as in A. Dashed line represents negative correlation. Significance in correlation: White (p<0.05) <Blue (p<0.005) <Green (p<0.0005) <Yellow (p<0.00001) as observed by Student’s t-test. (F) Heat maps for changes in gene expression downstream to insulin signaling in response to repeated insulin stimulation (N=2, n=3)

## Discussion

Coupling nutrient inputs to cellular metabolism, survival and growth is intrinsically dependent upon Insulin signalling (IS). Hypo- and hyper-activation of IS leads to various patho-physiologies including diabetes, accelerated aging and cancer, which are attributed to under- or over-phosphorylation of certain IS components (Arcidiacono et al., 2012; Guo, 2014; Hill and Milner, 1985; Shimobayashi et al., 2018; Shoelson et al., 2006; Vigneri et al., 2020). Despite this our ability to tweak the cascade to restore balance between metabolic and mitogenic arms has been limited by paucity of information vis-à-vis parameters that govern network topology. In this study, using mathematical and experimental approaches, we have provided fundamental insights into kinetic parameters that dictate emergent properties of IS and its architecture, under various physiological contexts.

Given the contribution of the liver in maintaining whole organismal physiology and insulin action, including development of metabolic diseases, we have specifically utilized primary hepatocytes for deciphering kinetic constants or determinants that exert a control over IS. It should be noted that while it is nearly impossible to recreate paradigms that mirror in-vivo conditions, we have employed insulin treatment regimens that mimic normo- and hyper-insulinemic states. Moreover, in-vivo complexity of insulin-dependent endocrine and paracrine networks would severely confound attempts to unveil kinetic determinants. Our study has revealed novel insights into kinetic control of insulin signalling and also provides a model to capture such parameters in other cells or tissue types, including by coupling other endocrine/paracrine inputs.

While genetic, biochemical and pharmacological perturbations have described inter-dependence of IS phosphorylation events, recent phospho-proteomic analyses have unravelled their temporal behaviour. However, the extent to which phosphorylation dynamics encode information as a function of insulin concentration and/or time is still unclear. For example, even though hypo-/hyper-phosphorylations at T^308^ and S^473^ are considered as proxy markers for AKT activity and signalling downstream to insulin, whether or not their kinetic differentials contribute to insulin responsiveness remains unknown. Here, we surprisingly found that while the gain and kinetics of pT^308^ (IR/PDK1 dependent) was independent of input strength, phosphorylation of S^473^ (downstream to mTORC2) correlated with change in ligand concentration and displayed highest gain in signal in response to a fed insulin input.

Further, in contrast to net gain in specific phosphorylation, dynamic range, which is undetermined for many signalling networks including IS, has been proposed to be a better predictor of cellular response. In this regard, our simulation and experimental data together revealed a large dynamic range for pAKT^T308^ and pAKT^S473^, which was nearly overlapping. This suggests that both these phosphorylations are equally responsive to relative change in insulin inputs vis-à-vis physiological fed-fast cycles wherein circulating concentrations vary between 0.1nM and 1.0 nM. Surprisingly, we also found that pT^308^ is a key determinant of network topology, which also highlights distinct properties of AKT phosphorylations in contributing to flow of information. Taken together these also raise the possibility of pT^308^ and pS^473^ acting as low pass and high pass filters with former being a permissive cue, which was hitherto unknown.

Noise in biology is generally regarded to be beneficial for regulating functional flexibility and has been well studied in the context of gene transcription. Given limited knowledge in this regard for signalling cascades (especially for IS), we checked for input versus variance in signal response for the nodal kinase AKT. It was interesting to note that noise in signaling was apparent at physiological concentrations of insulin (0.1-1 nM) while it was substantially diminished in hyper-insulinemic regimes. This hinted towards reduced flexibility in signaling under hyper-insulinemic states.

Signal stratification is crucial for sustenance of downstream information even upon input extinction. We discovered that signals are stratified with differential gating, in an insulin concentration dependent manner, with kinetic barriers/gates emerging at both low and hyper-insulinemic concentrations. These bring to the fore the need to address mechanisms that contribute to these kinetic barriers by affecting K_ON_/K_OFF_ ratios of phosphorylation events, in the future. We propose such components would be very attractive candidates for therapeutic interventions to regulate insulin signalling and maintain network properties.

In addition to differential kinetic gating, connectedness between signalling components determines topology of the network. Despite several studies on signalling cascades across biological systems, little is known about if/how these parameters contribute to topology, except in cases where simulations have been carried out for artificial signalling systems. Our iterative experimental-simulation approach has revealed that maximum connectivity between the signalling nodes, which is often used as a measure of network robustness, is achieved at physiological concentrations of insulin. Conversely, the network breaks at hyper-insulinemic states. Importantly, we also underscore the significance of each of the phosphorylations in maintaining the robustness of the topology under normo-insulinemic states.

Others and we have found that metabolic cues under fasting conditions elicit anticipatory molecular mechanisms to mount an efficient fed response (Chattopadhyay et al., 2020; Maniyadath et al., 2019; Shaw et al., 2020). Given that fasting insulin (0.1nM) is pulsatile with a frequency of 10-15min, our findings have shown that this rewires fed IS dynamics. Strikingly, we found that coupling low pulsatile inputs with 1.0nM insulin stimulation, as in the case of fasted to fed transition, enhanced net gain in phosphorylation of some (pAKT^T308 and S473^ and pGSK3β^S9^) but not all components, akin to memory or anticipation. Conversely, insulin resistance is associated with repeated insulin/nutrient inputs and hyperinsulinemia. Our study also describes kinetic changes in IS dynamics, which can be either causal or consequential to reduced sensitivity under these conditions. Notably, we found that repeated stimulation with fed concentrations of insulin damped the AKT response while upregulating pERK indicating a disbalance between metabolic and mitogenic arms. This is important because overactivation of either metabolic and/or the mitogenic arm has been described in literature as a driver of metabolic diseases and cancer (Altomare and Testa, 2005; Burotto et al., 2014; De Luca et al., 2012; Shaw and Cantley, 2006). Here, we would like to specifically highlight that the signaling network is most robust in response to fed insulin inputs, which is pulse primed by fasting insulin. Our findings posit that repeated and/or high insulin inputs, including in a clinical setting could lead to perturbed networks with possible pathological manifestations.

In conclusion, our results unravel hitherto unknown kinetic constraints that exert control over components of insulin signaling. Notably, we illustrate that these kinetic parameters are intrinsically linked to insulin concentrations as in normo- and hyper-insulinemic states. Given that a discordant signal flow between metabolic and growth-factor arms is associated with diseases, our findings provide fundamental insights into factors that govern this coupling. Our study also raises the possibility of impaired biological outputs in the context of therapeutic interventions using insulin, which have been largely guided by glycemic control. We highlight the importance of discovering novel regulatory parameters/nodes to complete our understanding of signaling cascades under both normal and pathological conditions.

## Materials and Methods

### Animals

2.5-3 month old C57BL/6NCr mice were used for hepatocyte isolation. The animals were housed under standard animal house conditions with a 12h day and night cycle. All procedures were done in accordance with the institute animal ethics committee (IAEC) guidelines.

### Primary Hepatocyte Isolation and culture

Male mice were sedated by giving intraperitoneal Thiopentone (Neon Laboratories Ltd., Mumbai, India) injection. Liver perfusion was done via inferior vena cava using 30mL Hank’s Balanced Salt Solution, HBSS (5.33 mM Potassium chloride, 0.44 mM KH_2_PO_4_, 4.16 mM NaHCO_3_, 137.93 mM NaCl, 0.338 mM Na_2_HPO_4_, pH 7.4) containing 5.5 mM Glucose (Sigma-Aldrich G8769), 25mM HEPES pH 7.2 (USB 16926) and 100mM EGTA (Sigma-Aldrich E3889). Hepatic portal vein was cut at one end in order to drain out the blood. Perfused liver was digested using collagenase (Sigma-Aldrich C5138) dissolved in 50mL Digestion medium {DMEM-LG (Sigma-Aldrich D5523), 15mM HEPES pH 7.2 (USB 16926) and Anti-Anti (Sigma-Aldrich A5955)}. Liver was cut into pieces, minced and incubated in Digestion Medium for 5min. The cells were strained using a 70µm cell strainer and centrifuged at 50G for 5min. Cell pellet was washed twice with DMEM-HG (Sigma-Aldrich D7777) and re-suspended in DMEM-HG containing 10% FBS (Gibco 16000044) for plating. Trypan blue staining was done to check cell viability. Cells were plated at a density of 7.5 × 10^5^ cells/60mm plate in collagen (Sigma-Aldrich C3867) coated plates (5µg/cm^2^). Cells were grown at 37°C and 5% CO_2_. Medium was changed 6h post plating to ensure proper cell adherence.

### Insulin Treatments

24h post plating, the hepatocyte medium was changed to 5% FBS containing DMEM-HG for 11h. Medium was changed to Earle’s Balanced Salt Solution, EBSS (Sigma-Aldrich E2888) for 6h to get a baseline (0m) signal. For one step insulin stimulation experiments, 0.1-100nM Insulin (Sigma-Aldrich I0516) in DMEM-HG was added to the hepatocytes and cells were collected at time points as described in the results. For pulsatile insulin treatments and repeated insulin stimulation, paradigm modifications are mentioned in Figure 4A and 5A. Every media change was preceded with a PBS wash to remove residual contamination.

### Protein Lysate Preparation

Hepatocytes were lysed in RIPA lysis buffer (50mM Tris pH 8.0, 150mM NaCl, 0.1% SDS, 0.5% Sodium deoxycholate, 1% Triton X-100, 0.1% SDS, 1 mM PMSF, Protease inhibitor cocktail and phosphatase inhibitor-Sigma-Roche 4906845001) for 30min. Cell debris were pelleted by centrifuging at 12,000rpm at 4°C for 15min. BCA assay kit (Sigma-Aldrich 9643) was used for protein estimation. Protein samples were boiled in a loading buffer (8% SDS, 40% glycerol, 240mM Tris pH 6.8, 0.2g bromophenol blue and 3.05g DTT) and stored at × 20°C.

### Western Blotting

50µg of protein was loaded onto SDS gel and run at 90V for stacking and 120V for resolving. Gels were transferred to ethanol activated PVDF membranes (Merck IPVH00010-IN) at 90V for 2h. Protein blotted membranes were blocked in 5% skimmed milk. Blots were cut according to protein molecular weight as indicated by pre-stained ladder (Abcam ab116028) and incubated overnight with the respective primary antibodies: AKT (CST 9272), pAKT^S473^ (CST 4060), pAKT^T308^ (CST 13038), ERK1/2 (CST 4695), pERK1/2^T202/Y204^ (CST 4376), GSK3β (CST 12456), pGSK3β^S9^ (CST 5558), pS6K^T389^ (CST 9234) and S6K (CST 2708). Blots were incubated with appropriate secondary antibodies (Anti-Rabbit IgG-Peroxidase Sigma-Aldrich A0545 and Anti-Mouse IgG-Peroxidase Sigma-Aldrich A9044) and imaged using GE Amersham Imager 600.

### RNA extraction, cDNA synthesis and RT-PCR

RNA extraction, cDNA synthesis and real time PCR was performed as per manufacturer’s instructions. Briefly, total RNA was extracted from hepatocytes using TRIzol reagent (Ambion-Invitrogen 15596-018) and 1ug of RNA was used to make cDNA using SuperScript IV RT Kit (Invitrogen 18090010). Quantitative PCR was done using KAPA SYBR® FAST Universal 2X qPCR Master Mix (KAPA Biosystems KK4601) and LightCycler 96 instrument (Roche). The list of primers used are depicted in the table below:

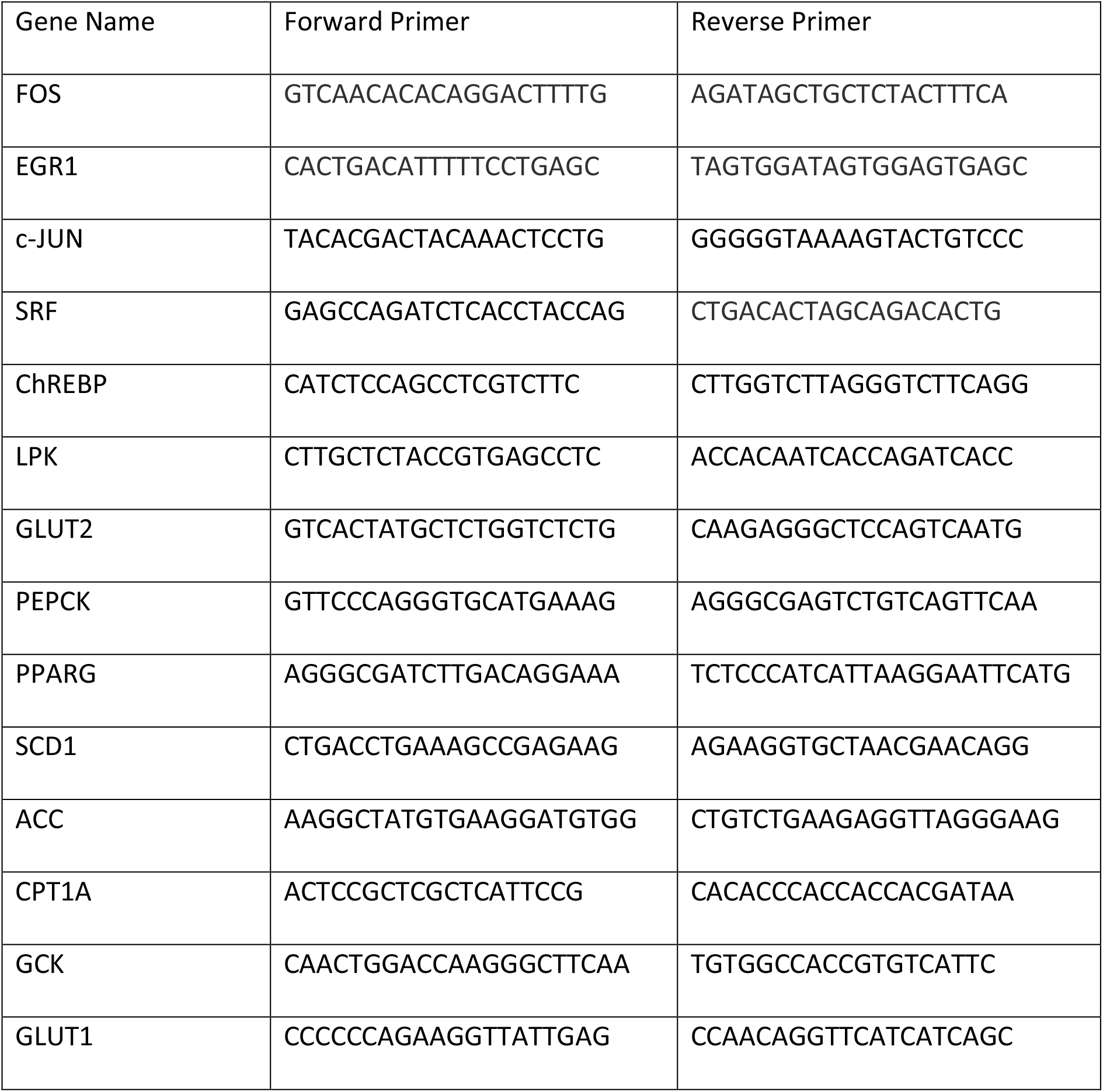
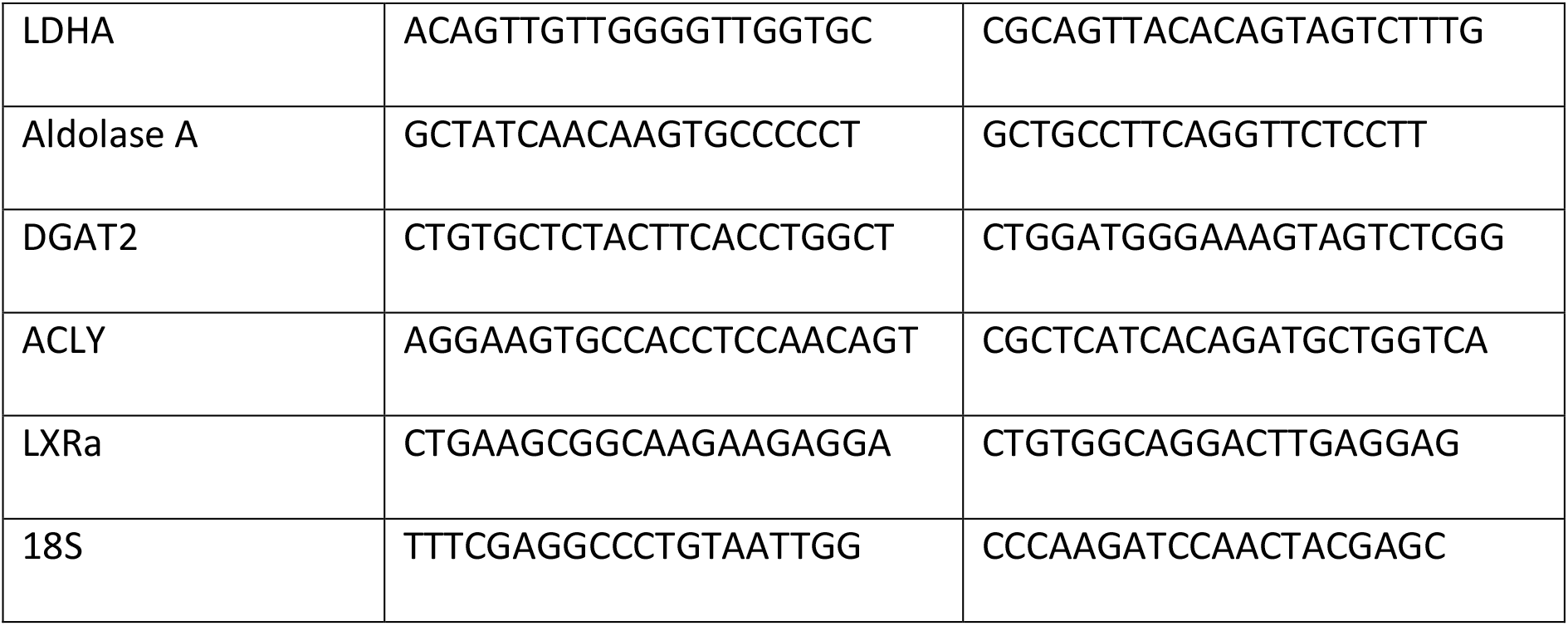

### Data Processing

Intensity measurements from the blots were done using Fiji-ImageJ software with corresponding background correction.

### Network Analysis

Network construction, visualization and analysis was performed using Cytoscape (version 3.7.2) using Pearson correlation data obtained from GraphPad Prism (version 8).

### Quantitation and Statistical Analysis

Data are expressed as means ± standard error of means (SEM). Statistical analyses were performed using Microsoft Excel (2013) and GraphPad Prism (version 8). Statistical significance was determined by the Student’s t test. A value of p ≤ 0.05 was considered statistically significant. *p ≤ 0.05; **p ≤ 0.01; ***p ≤ 0.001.

### Calculation of z-score and true/false discovery rate (TDR/FDR)

Z-score was calculated by computing the difference between experimental and simulation for each time point. Like in a paired t-test or z-test, we computed the z-value by calculating the mean of the differences and dividing by the standard deviation. If z < 1.96, the null hypothesis — that the experimental and simulation mean values are statistically the same was accepted. For computing FDR, difference between experimental and simulation data was divided by standard errors for different insulin concentrations and z-values were calculated. z > 1.96 it is was rejected and counted as false discovery and further averaged to obtain the false discovery rate. For two different insulin concentrations, we define a true discovery rate, which is calculated by taking difference between mean values and dividing by root of sum of square of standard errors. If z > 1.96, the null hypothesis is rejected and counted as true discovery and averaged to obtain the true discovery rate.

### Estimation of Parameters for Deterministic Simulations

The proposed pathway of insulin signaling is modelled in terms of ordinary differential equations by using mass action kinetics (Alon, 2019; Klipp et al., 2016). The set of equations that is used to define the insulin signaling network, which is considered both single (denoted by ‘p’) and double phosphorylation (denoted by ‘pp’) events and feedback inhibitions, are depicted in Figure S3. We have solved these sets of ordinary differential equations using MatLab version R2015b, Math Works. The parameters of the model (rate constants of reactions and initial amounts of proteins) were decided such that the model fitted with the experimental data. The constrained nonlinear optimization technique was implemented using the ‘fmincon’ function in MatLab to provide parameters that fit best with the available data. That is, we optimized (minimized) the function:

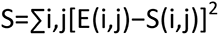

where, E(*i, j*) and S(*i*, *j*) are the experimental and simulation data, respectively, for the *i*^th^ protein component at *j*^th^ time point. The function essentially measures the deviation of the experimental data, and is defined as the sum of the squares of the differences between experimental measurements and simulated trajectories over all the measured proteins. We took 100 independent runs of the program to estimate these parameters. We choose the parameters that correspond to a minimum objective function value resulting in a good fit. All parameters were obtained for different insulin concentrations.

### Parameters used in Stochastic Simulations

This reaction network is simulated by using the kinetic Monte Carlo based Doob-Gillespie Algorithm (Doob, 1942, 1945; Gillespie, 1976, 1977). The rate constants of respective reactions are taken from the deterministic model proposed earlier. These rate constants are converted to stochastic framework by using appropriate conversion factors. For unimolecular reactions, the deterministic rate constants (k_j_) and stochastic rate constants (c_j_) are numerically equal. For bimolecular reactions, when two reactants correspond to different proteins, the stochastic rate constants (c_j_) are equal to k_j_/V, where V is the system volume (Gillespie, 1976, 1977). The concentration of proteins from a deterministic regime are converted to the number of proteins per cell by multiplying the concentrations by N_a_ × V, where N_a_ is the Avogadro’s number and V = 3×10^−12^ liters is the estimated volume of a cell. During the simulation, in any particular iteration from a given reaction network a single bio-chemical reaction and the subsequent time step is chosen randomly. In this way, a single stochastic trajectory is generated by running the simulation for a desired time. Further, many more realizations of this trajectory are generated to compute different moments (e.g. mean, standard deviation) of the probability distributions.

Estimation of Insulin Molecules: To estimate the number of insulin molecules from concentration, we assumed a spherical shell (around the cell membrane) of 20nm size and computed the corresponding volume. Assuming a spherical cell of volume V = 3×10^−12^ litres the volume of this shell is ΔV = 0.0199×10^−12^ litres. Hence the number of insulin molecules are estimated by calculating N_a_ × ΔV, where N_a_ is the Avogadro’s number.

i. For 0.1 nM Insulin: the number of insulin molecules are found to be 1.1985 molecules per spherical shell (1 insulin per spherical shell)
ii. For 1 nM Insulin: the number of insulin molecules are found to be 11.9857 molecules per spherical shell (12 insulin molecules per spherical shell)
iii. For 10 nM Insulin: the number of insulin molecules are found to be 119.8577 molecules per spherical shell (120 insulin molecules per spherical shell)

### Calculation of decay rate

Decay times are calculated for dynamic protein concentrations measured and simulated in our studies by fitting an exponential function (e^-kt^) from the time point at which peak intensity is maximum to the final time point at which the intensity falls down.

### Calculation of parameters in kinetic gating

The kinetic gating in the signaling cascade is studied by taking the ratio of phosphorylation rate constant (K_ON_) and the de-phosphorylation rate constant (K_OFF_) of each biochemical reaction. In case of the degradation reactions, their rate constants are incorporated by averaging with the rate constants of appropriate reactions.

## Acknowledgements

We acknowledge Dr. Shital Suryavanshi, Dr. Kalidas Kohale and TIFR-Animal House staff for help with animal breeding and maintenance. This research has been supported by funds to U.K.-S. {DAE-TIFR (Government of India grant - 12P0122), Department of Biotechnology (DBT, India grant BT/PR4972/AGR/36/714/2012) and Swarnajayanti fellowship (DST Government of India grant DST/SJF/LSA-02/2012-13)}, R.P. (DBT, India grant BT/HRD/NBA/39/12/2018-19) and S.K. (Department of Science and Technology, India under Science and Engineering Research Board (SERB) - National Post-doctoral Fellowship (NPDF) with file number: PDF/2017/002502). We also thank the members of UK lab, TIFR for useful discussions and constructive comments.

## Competing interests

The authors declare that no competing interests exist.

---

**Figure 1 – figure supplement 1:**
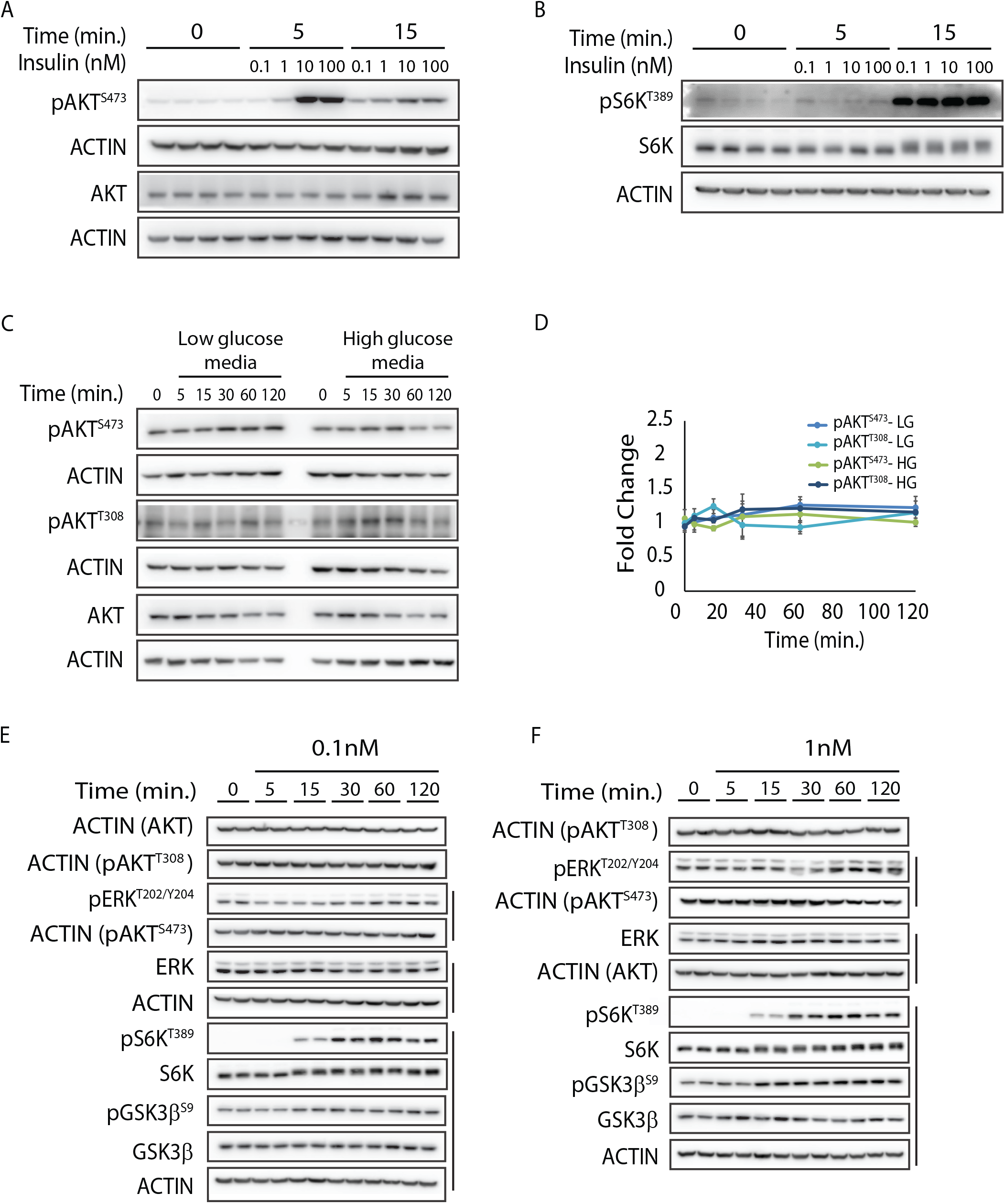
(A-B) Normalization controls to correct for baseline signal. Representative samples were loaded to ensure that zero-minute time points across experiments were similar. (B) was over-exposed to get signal at zero minute. (C) Control for experimental paradigm to score for insulin inputs. Treatment with only high/low glucose and amino acid containing culture medium does not activate AKT signaling and indicates that the changes in phosphorylation are insulin dependent. (D) Quantitation for temporal changes in phosphorylations for C. (E and F) Representative blots for levels of pERK^T202/Y204^, pS6K^T389^ and pGSK3β^S9^ following insulin stimulation of 0.1 and 1 nM, as indicated. Respective total proteins and actin were used for normalization

**Figure 1 – figure supplement 2.**
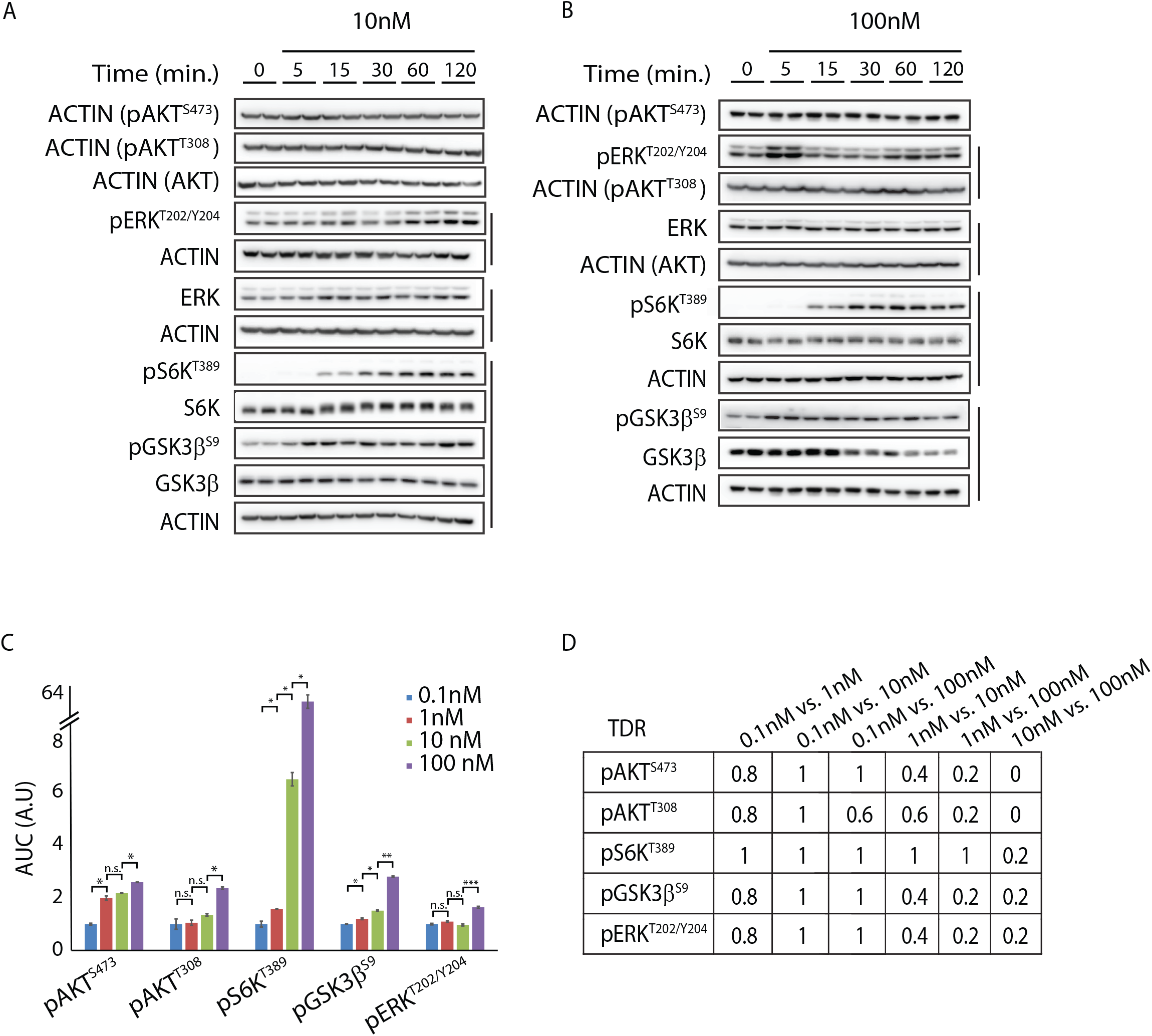
(A and B) Representative blots for levels of pERK^T202/Y204^, pS6K^T389^ and pGSK3β^S9^ following insulin stimulation of 10 and 100 nM, as indicated. Respective total proteins and actin were used for normalization. (C) Comparison of area under the curve (AUC) with increasing insulin concentration. Asterisk depicts p values (*p<0.05, **p<0.005 and ***p<0.0005) as observed by Student’s t-test. (D) True discovery rates computed across insulin concentrations, see methods.

**Figure 1 – figure supplement 3.**
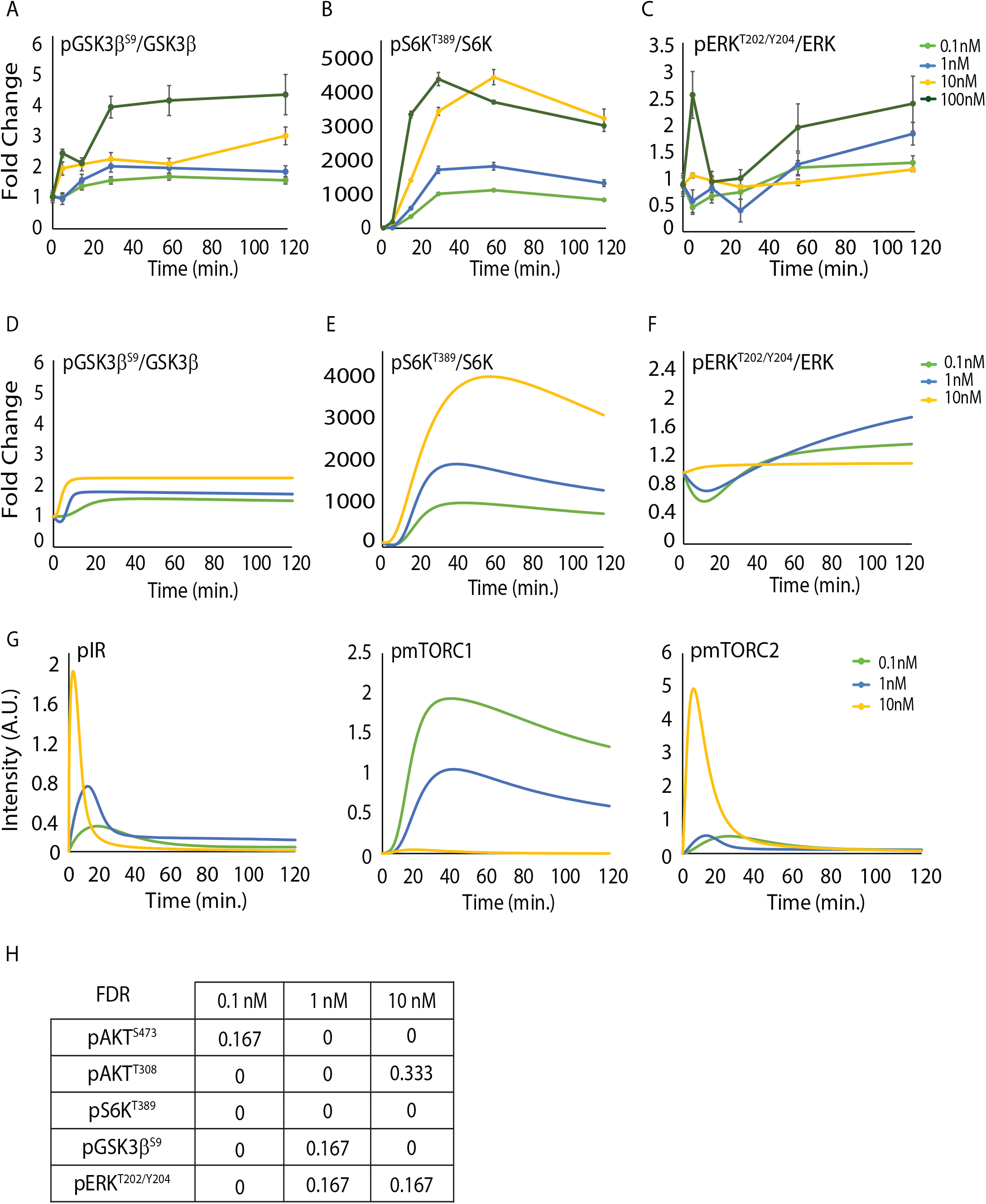
(A-C) Quantitation for temporal changes in pGSK3β^S9^(A), pS6K^T389^ (B) and pERK^T202/Y204^ (C) from experimental data shown in Figure 1 – figure supplement 1E-F and 2A-B. Fold changes for each concentration are with respective to their own 0m time point. Data presented is mean ± s.e.m. (N=4, n=4). (D-F) Quantitation for temporal changes in pGSK3β^S9^ (D), pS6K^T389^ (E) and pERK^T202/Y204^ (F) from mathematical simulations using differential equations. (G) Simulated temporal changes in pIR, pmTORC1 and pmTORC2 following insulin stimulation, as indicated. (H) False discovery rates giving degree of concordance between simulated and experimental data, see methods

**Figure 2 – figure supplement 1.**
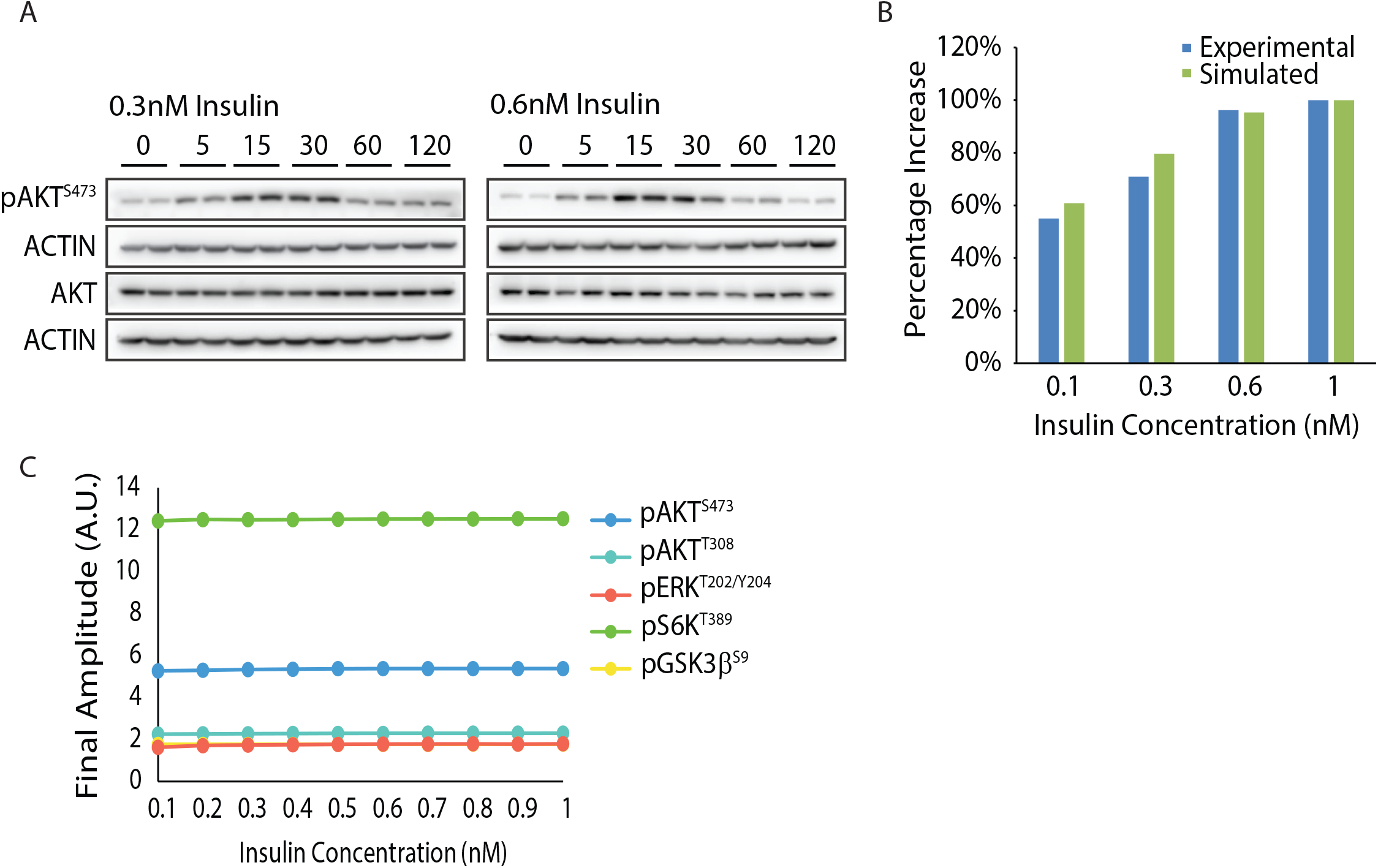
(A) Representative blots for level of pAKT^S473^ kinetics under insulin concentrations of 0.3 and 0.6 nM. Total AKT and actin were used for normalization. (B) Concordance between experimental and simulated data for extent of phosphorylation at pAKT^S473^ following stimulation by intermediate insulin concentrations. (C) Estimated final amplitude for signaling components at 120m as a function of varying insulin concentrations

**Figure 3 – figure supplement 1.**
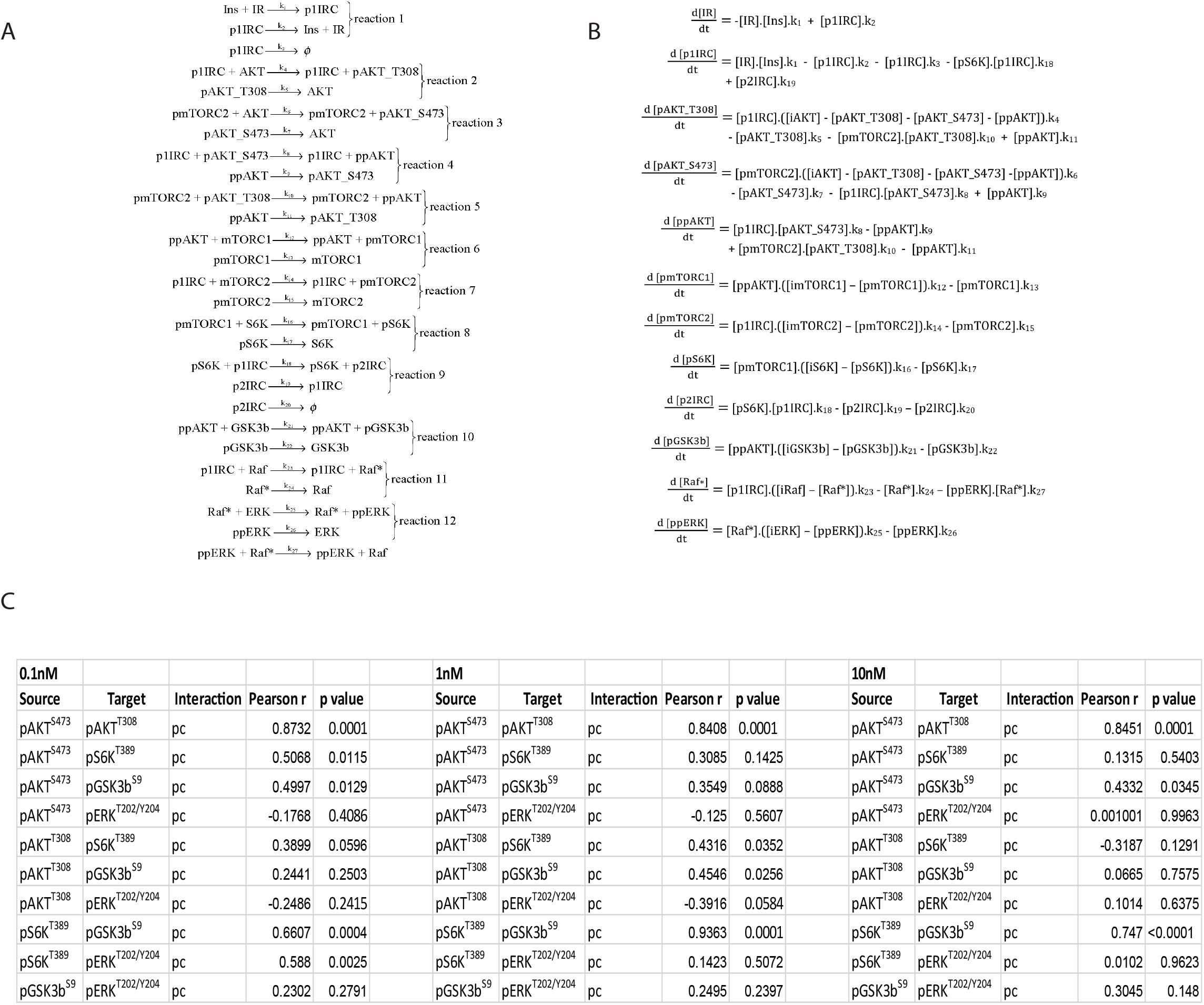
(A) Reactions corresponding to numbers on y-axis in Figure 3A. (B) Ordinary differential equations corresponding to reactions mentioned in A. (C) Pearson r and corresponding p values used for computing networks in Figure 3D. and Figure 3- figure supplement 2

**Figure 3 – figure supplement 2.**
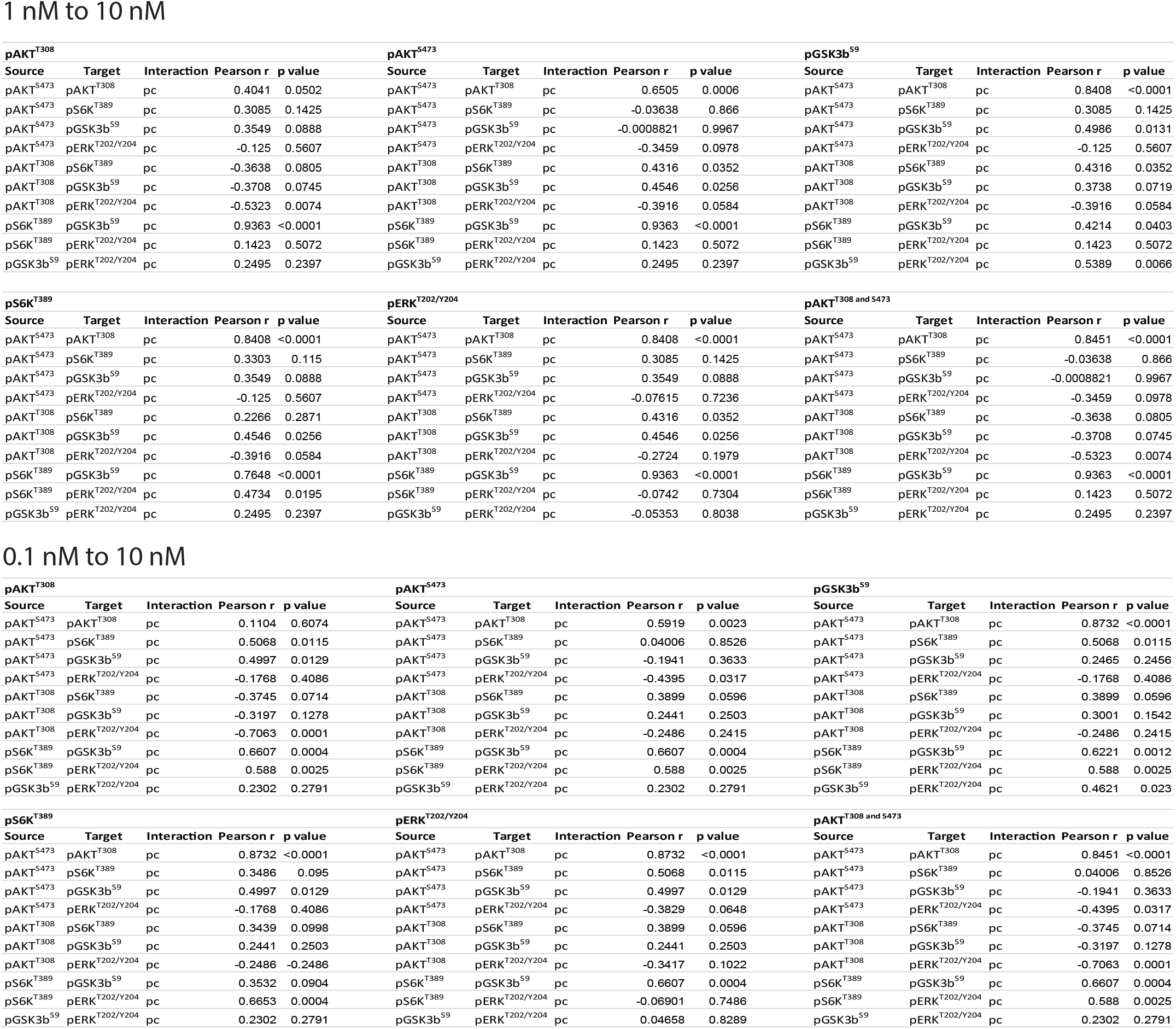
Pearson r and corresponding p values used for computing networks in Figure 3F and Figure 3- figure supplement 3

**Figure 3 – figure supplement 3.**
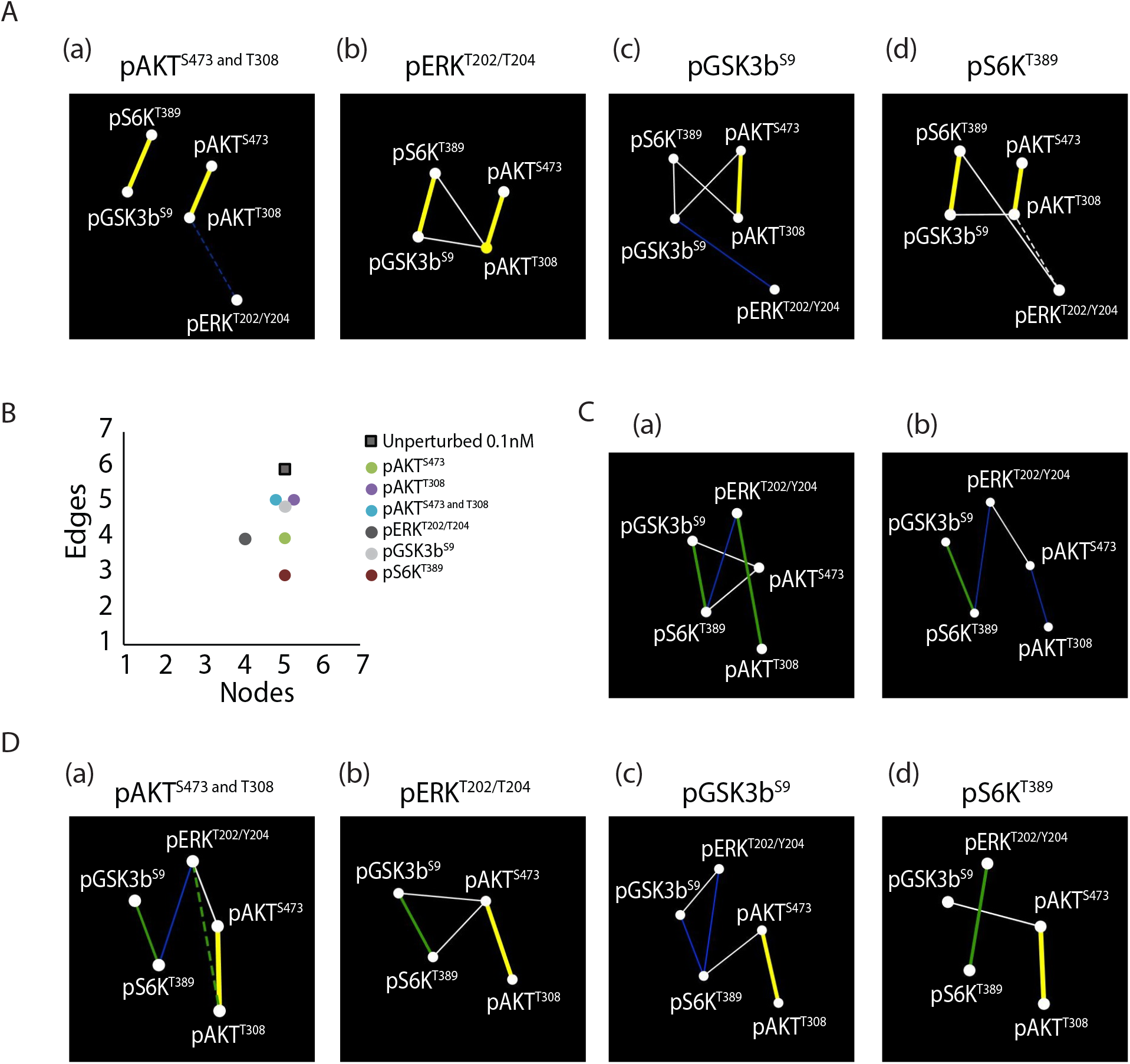
(A) Network maps of 1 nM insulin perturbed with 10 nM pAKT^T308 and S473^ (a), pERK^T202/Y204^ (b), pGSK3b^S9^ (c) and pS6K^T389^ (d). (B) Number of edges and nodes in a 0.1 nM network substituted with 10 nM values, as indicated. (C) Network maps of 0.1 nM insulin perturbed with 10 nM pAKT^T308^ (a) and pAKT^S473^ (b). (D) Network maps of 0.1 nM insulin perturbed with 10 nM pAKT^T308 and S473^ (a), pERK^T202/Y204^ (b), pGSK3b^S9^ (c) and pS6K^T389^ (d).

**Figure 4 – figure supplement 1.**
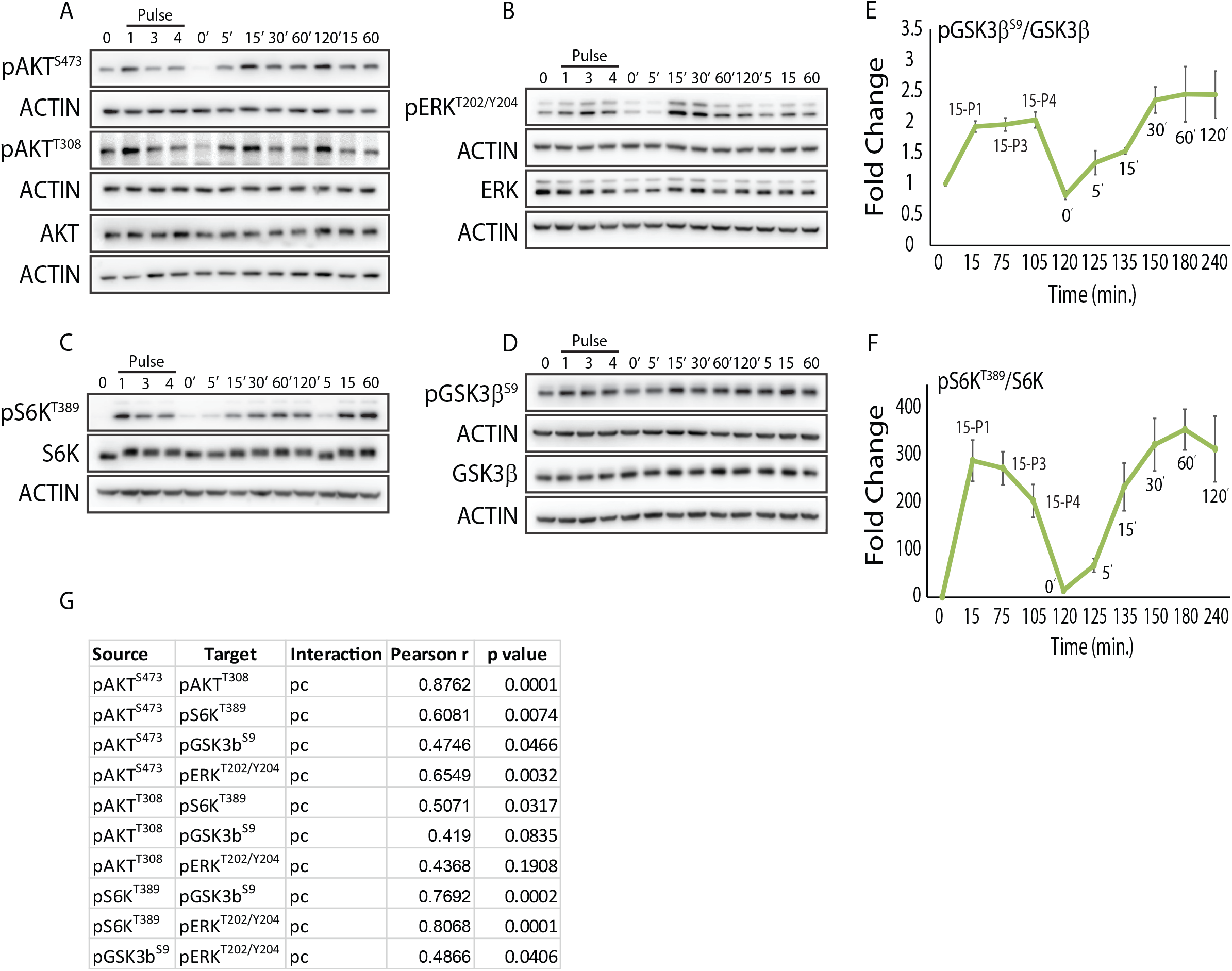
(A-D) Representative blots for levels of pAKT^T308^ (A), pAKT^S473^ (A), pERK^T202/Y204^ (B), pS6K^T389^ (C) and pGSK3β^S9^ (D) following pulsatile insulin stimulation, as indicated in Figure 4A. Respective total proteins and actin were used for normalization kinetics. (E and F) Quantitation for temporal changes in phosphorylations at pGSK3β^S9^ (E) and pS6K^T389^ (F) following insulin pulses. Fold changes for each concentration are with respective to their own 0m time point. Data presented is mean ± s.e.m. (N=4)

**Figure 5 – figure supplement 1.**
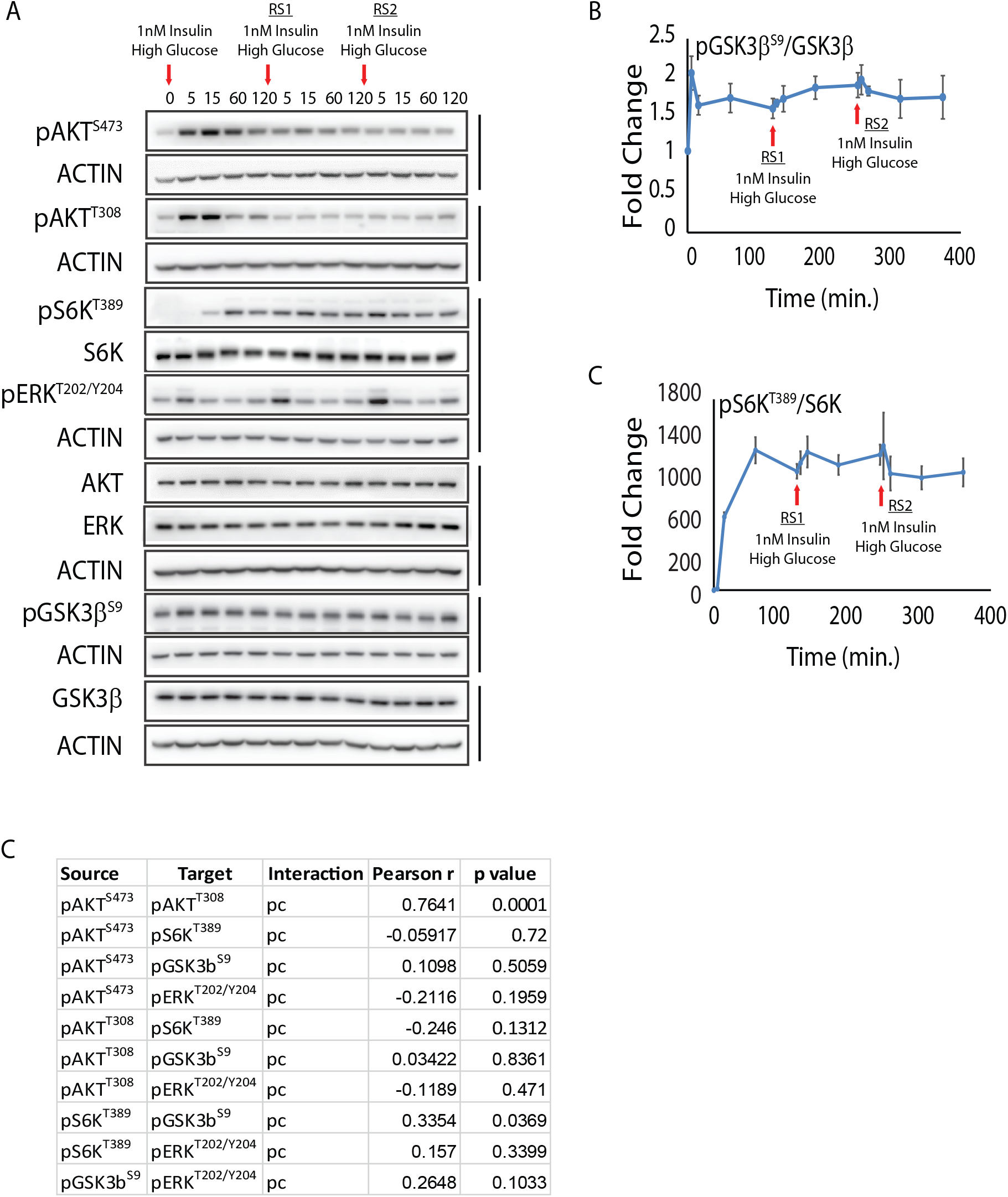
(A-D) Representative blots for levels of pAKT^T308^ (A), pAKT^S473^ (A), pERK^T202/Y204^ (B), pS6K^T389^ (C) and pGSK3β^S9^ (D) following repeated insulin stimulation, as indicated in Figure 5A. Respective total proteins and actin were used for normalization kinetics. (E and F) Quantitation for temporal changes in phosphorylations at pGSK3β^S9^ (E) and pS6K^T389^ (F) following repeated insulin stimulation. Fold changes for each concentration are with respective to their own 0m time point. Data presented is mean ± s.e.m. (N=4).

## Source Data

**Figure 1 – source data 1:** Blot intensity and quantitation for pAKT^S473^ and pAKT^T308^

**Figure 1 – source data 2:** Values for pAKT^S473^ and pAKT^T308^ from deterministic simulations used for plotting Figure 1F-G

**Figure 1 – source data 3:** Values for pAKT^S473^ from stochastic simulations used for plotting Figure 1H

**Figure 1 – source data 4:** z-score computation for concordance between simulated and experimental data

**Figure 1 – figure supplement 1 – source data 1:** Blot intensity and quantitation for Figure 1 - figure supplement 1C-D

**Figure 1 – figure supplement 2 – source data 1:** Calculations for computing area under the curve

**Figure 1 – figure supplement 2 – source data 2:** TDR computation for comparison across insulin concentration

**Figure 1 – figure supplement 3 – source data 1:** Blot intensity and quantitation for pGSK3β^S9^, pS6K^T389^ and pERK^T202/Y204^ used for plotting Figure 1 – figure supplement 3A-C

**Figure 1 – figure supplement 3 – source data 2:** Values for pGSK3β^S9^, pS6K^T389^ and pERK^T202/Y204^ from deterministic simulations used for plotting Figure 1 – figure supplement 3D-F

**Figure 1 – figure supplement 3 – source data 3:** Values for pIR, pmTORC1 and pmTORC2 from deterministic simulations used for plotting Figure 1 – figure supplement 3G

**Figure 1 – figure supplement 3 – source data 4:** FDR calculations for degree of concordance between simulated and experimental data

**Figure 2 – source data 1:** Percentage gain computation w.r.t. Figure 2A

**Figure 2 – source data 2:** Values for peak intensity and decay time used for plotting Figure 2B

**Figure 2 – source data 3:** Values for peak intensity used to plot Figure 2C

**Figure 2 – source data 4:** Computation of dynamic range for Figure 2D

**Figure 2 – figure supplement 1 – source data 1:** Blot intensity and quantitation for extent of phosphorylation at pAKT^S473^ following stimulation by intermediate insulin concentrations in Figure 2- figure supplement 1A-B

**Figure 2 – figure supplement 1 – source data 2:** Values for final intensity at 120min. used to plot Figure 2- figure supplement 1C

**Figure 3 – source data 1:** K_ON_/K_OFF_ used for plotting Figure 3A

**Figure 3 – source data 2:** Coefficient of variation used for plotting Figure 3B

**Figure 3 – source data 3:** Computations for Pearson r and statistical significance used to construct Figure 3C

**Figure 3 – source data 4:** Values for edges and nodes corresponding to Figure 3E and Figure3- figure supplement 3B

**Figure 3 – figure supplement 1 – source data 1:** Rate constants corresponding to equations in Figure 3-figure supplement

**Figure 4 – source data 1:** Blot intensity and quantitation for extent of phosphorylation at pAKT^S473^, pAKT^T308^ and pERK^T202/Y204^ corresponding to Figure 4B-D

**Figure 4 – source data 2:** Computations for Pearson r and statistical significance used to construct Figure 4E

**Figure 4 – source data 3:** Ct values for changes in gene expression corresponding to Figure 4F

**Figure 4 – figure supplement 1 – source data 1:** Blot intensity and quantitation for extent of phosphorylation at pGSK3β^S9^ and pS6K^T389^ corresponding to Figure 4- figure supplement 1E-F

**Figure 5 – source data 1:** Blot intensity and quantitation for extent of phosphorylation at pAKT^S473^, pAKT^T308^ and pERK^T202/Y204^ corresponding to Figure 5B-D

**Figure 5 – source data 2:** Computations for Pearson r and statistical significance used to construct Figure 5E

**Figure 5 – source data 3:** Ct values for changes in gene expression corresponding to Figure 5F

**Figure 5 – figure supplement 1 – source data 1:** Blot intensity and quantitation for extent of phosphorylation at pGSK3β^S9^ and pS6K^T389^ corresponding to Figure 5- figure supplement 1B-C

